# Cross-frequency routing in a hippocampo–cortical circuit during probabilistic reversal learning

**DOI:** 10.64898/2025.12.16.694625

**Authors:** Alejandro Aguilera, Nelson Espinosa, Mauricio Caneo, Guillermo Lazcano, Ariel Lara-Vasquez, Pablo Fuentealba

## Abstract

Learning under uncertainty requires detecting latent changes in environmental contingencies and flexibly adjusting choice strategy. To model this process and identify associated circuit dynamics, we trained rats on a two-armed bandit task with uncued reward reversals while simultaneously recording local field potentials from the circuit comprised by dorsal hippocampus (CA1d), lateral entorhinal cortex (LEC), and prefrontal cortex (PFC). Behavioral performance improved with training and progressively shifted from outcome-reactive exploration to an exploitation-biased strategy, quantified using a model-free index. Mixed-effects modeling combining behavioral and neural metrics identified tonic cortical synchrony in theta and fast-gamma bands as a negative session-scale performance marker. Theta and gamma oscillations were coordinated, as cross-regional phase–amplitude coupling showed that CA1d theta phase gated fast-gamma activity in PFC, while biasing both slow- and fast-gamma bursts in LEC, which selectively increased at goal-reaching. Neuronal spiking showed cross-regional phase-locking to both theta and gamma rhythms, indicating synchronized timing across the hippocampo-cortical circuit. Finally, distributed neuronal spiking across the circuit represented goal-approach, yet only prefrontal neurons ramped during goal-reaching, suggesting a role in outcome assessment, while hippocampal and entorhinal units transiently suppressed, consistent with locomotor tracking and resetting. These results reveal a tonic cortical connectivity marker of ongoing performance during reversal learning and dissociate it from hippocampal theta mechanisms that selectively organize cortical gamma bursts and spike timing. Together, these results provide mechanistic insight into how frequency-specific hippocampo-cortical interactions dynamically reconfigure to support strategy transitions, offering a circuit-level framework for understanding adaptive decision-making under uncertainty.

**Significance Statement:** Animals often learn in uncertain environments, where they must decide whether to keep exploiting a known option or explore alternatives. We trained rats on a probabilistic choice task while recording brain activity from hippocampus, entorhinal cortex, and prefrontal cortex. We found that performance improvements were linked to an ongoing interaction between entorhinal and prefrontal areas, whereas hippocampal signals changed mainly with training and controlled the timing of brief, fast activity bursts in cortex. Near reward, prefrontal neurons ramped up while hippocampal and entorhinal neurons were suppressed. Together, these results separate brain signals that track current performance from those that shape cortical dynamics during learning.

## Introduction

Effective decision-making under uncertainty requires the ability to detect hidden changes in the environment and update behavioral strategies accordingly (1, 2). This challenge is exemplified by probabilistic reversal learning tasks, such as the two-armed bandit paradigm (3, 4), in which agents must continuously arbitrate between exploring alternative options and exploiting known rewards as reward contingencies shift unpredictably (5, 6). Resolving this exploration–exploitation tradeoff is central to flexible behavior, and uncovering the circuit-level mechanisms that support it remains a major subject in systems neuroscience (7–9).

Converging evidence implicates a distributed hippocampo–cortical network, centered on the prefrontal cortex (PFC), in guiding adaptive strategy selection (10–12). Within this circuit, the dorsal CA1 (CA1d) region of the hippocampus, the lateral entorhinal cortex (LEC), and medial PFC form an interconnected circuit that integrates contextual, mnemonic, and executive information to guide decision-making. Indeed, PFC maintains internal representations of latent task states and supports behavioral regulation under uncertainty (13–15). CA1d provides contextual information and supports latent state inference, particularly when external cues are ambiguous (16, 17). LEC, in turn, encodes local option identity and recent outcome history, delivering structured content to both hippocampus and cortex (18–20). These regions are anatomically linked through bidirectional projections, forming a closed-loop anatomical substrate for iterative learning and credit assignment (21–26).

A prevailing hypothesis is that frequency-specific oscillations coordinate communication within this circuit. Theta rhythms (4–10 Hz) align excitability across distant regions and support temporal structuring of information flow, while gamma rhythms (30–100 Hz) convey local computations within and between regions (27–29). Cross-frequency interactions, such as theta-phase/gamma-amplitude coupling, may provide a mechanism for routing information dynamically across nodes of the network depending on task demands (30). Prior work has shown increased theta synchrony between hippocampus and PFC during working memory, spatial navigation, and memory-guided decision-making (31–33), but it remains unclear how these dynamics support learning in uncertain environments.

Here, we address this question by recording local field potentials and unit activity simultaneously from CA1d, LEC, and PFC as rats learned a self-paced two-armed bandit task with uncued reversals. We quantified strategy at the session-scale using a model-free index derived from simple behavioral patterns following wins and losses, and used linear mixed models to identify neurophysiological metrics associated with task performance and learning stage (34, 35).

We identified novel neural dynamics underlying decision-making and strategy adaptation. Training induced a shift from outcome-reactive exploration to an exploitation-biased strategy, which was linked to changes in cortico-cortical connectivity, particularly LEC–PFC theta–gamma coupling. Moreover, hippocampal theta oscillations, particularly in CA1d, structured cortical gamma bursts and spike timing across the circuit, supporting the organization of temporal windows for processing. These results provide insights into how hippocampo–cortical circuits dynamically reconfigure to support adaptive decision-making under uncertainty. This framework offers insight into how distributed systems communicate to resolve uncertainty and adapt behavior, and helps to explain the functional role of theta–gamma interactions in mesoscale coordination.

## Results

### Training-dependent strategy shift in a probabilistic reversal learning task

We trained rats on a two-armed bandit task, a commonly used probabilistic serial reversal learning task (5, 6), in which animals made self-directed binary choices to maximize reward collection across trials (**Fig. 1A**). Within each block (27-30 trials), reward probabilities were fixed for the two options (0.83:0.12), and blocks switched without explicit cues; because outcomes were stochastic, transitions could only be inferred by trial-and-error (36). Across sessions, reward rate increased with training (P = 5.3×10^-10^; **Fig. 1B**). This effect was not the result of changes in opportunity or locomotion, since adding trials per session and mean running speed as covariates left the training effect significant (Table S1).

**Figure 1.**
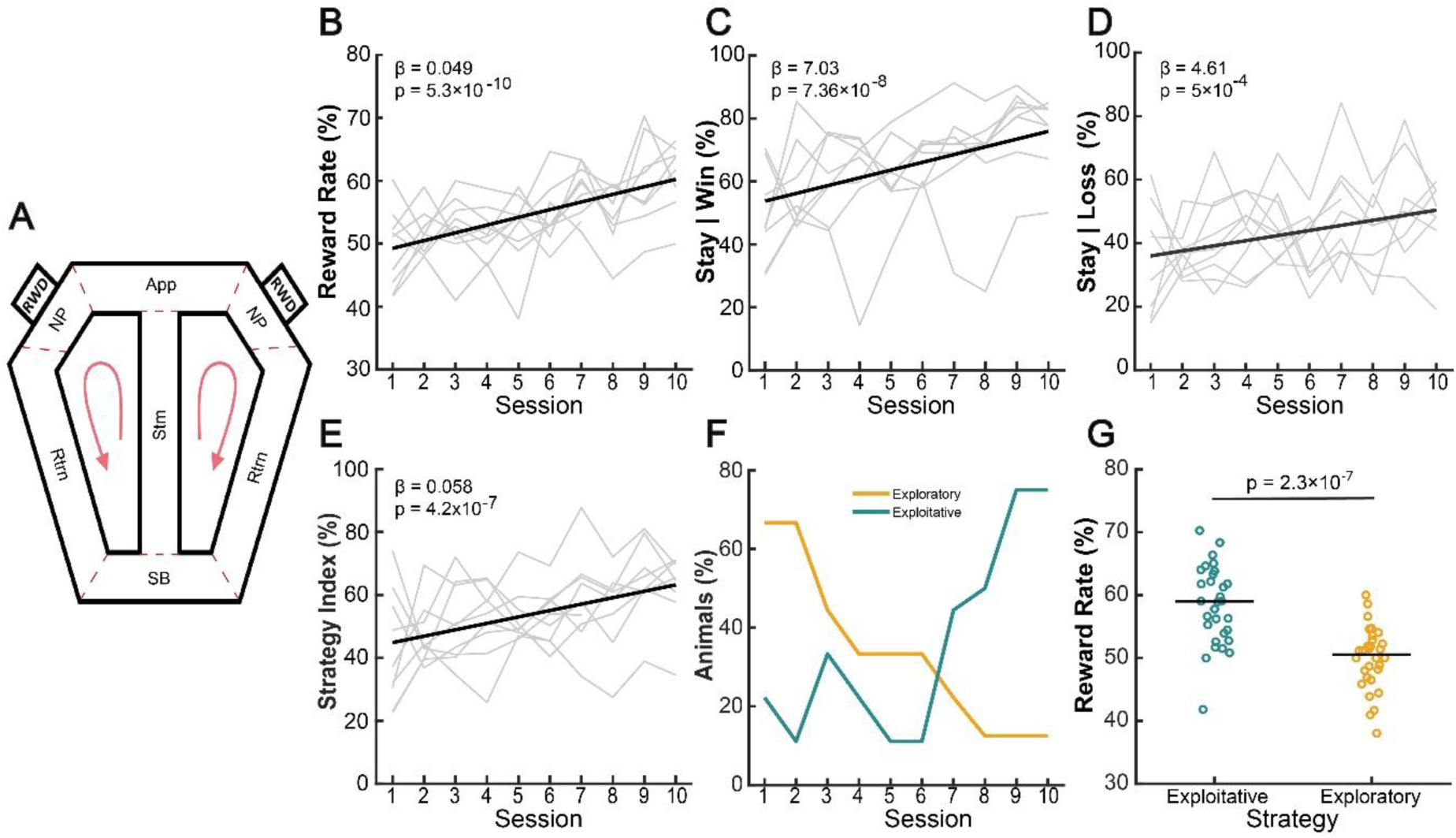
Learning and strategy during probabilistic reversal learning. **A,** Schematic of the task apparatus and behavioral epochs (red dashed boundaries): Start box (SB) → Stem (Stm) → Approach (App) → Nose-poke (NP) → Return (Rtrn). Animals ran in a fixed direction (red arrows) and chose an arm on each lap; the currently rewarded side (rwd) changed across unsignaled blocks. **B,** Session-wise reward rate (rewarded trials / valid trials; valid = total trial duration ≤ 100 s). A binomial GLMM (logit link; random intercepts and session slopes by rat) showed a positive effect of session (β = +0.049, P = 5.3×10⁻¹⁰), indicating robust improvement across training sessions. Thin gray lines, individual rats; black line, model population prediction. **C,** Win-stay probability increased with training (mixed-effects regression on session values: β = +7.03 percentage points per session, P = 7.36×10⁻⁸), consistent with growing exploitation of rewarded actions. **D,** Lose-stay increased over training (β = +4.61, P = 5.26×10⁻⁴), indicating that animals became more likely to maintain their choice despite isolated errors. Thin gray lines show individual animals; thick line indicates the fixed-effects prediction. **E,** Strategy Index (SI) per rat and session (valid trials only). S = 0.5×(SW + SL), where SW is stay|win and SL is stay|loss (persistence after a single loss). A linear mixed-effects model revealed an increase with session (β = +0.058, P = 4.2×10⁻⁷). Gray lines, individual rats; black line, model population prediction. **F,** Fraction of animals classified as Exploratory or Exploitative on each session based on predefined criteria; Mixed sessions are omitted from the plot but counted in the daily denominator. **G,** Session-wise reward rate by strategy. Dots, individual sessions; black lines, group medians. Exploitative sessions achieved higher reward rates than exploratory sessions (59.0% vs 50.6%; mixed-effects strategy effect: P = 2.3×10⁻⁷).

We next assessed behavioural strategies during decision-making in individual training. With training, rats became more likely to repeat a choice after a reward (win-stay; **Fig. 1C**) and more tolerant to a single loss (lose-stay; **Fig. 1D**). In parallel, both lose-shift and win-shift declined steadily (Fig. S1). To summarize contingencies in a model-free way, we defined a simple Strategy Index (SI) from stay-win*stay-lose behavior (Fig. S2). SI quantifies repeating after wins while tolerating isolated losses (**Fig. 1E**). Using data-driven thresholds we classified individual sessions (exploitative: SI > 0.60 and P(stay|win) > 0.70; exploratory: SI < 0.50 and P(stay|win) < 0.60; mixed otherwise), resulting in a relatively uniform distribution, with exploitative 38.96% (30/77), exploratory 38.96% (30/77), and mixed 22.08% (27/77) (Fig. S2). When assessing the temporal evolution of strategies, we found that the prevalence of exploitative sessions increased with training while exploratory sessions declined in parallel (**Fig. 1F**). We then modeled behavioral performance with a generalized linear mixed model predicting session reward rate from training and strategy (Table S2). We confirmed that reward rate increased with training within rats (β = 0.126, P = 3.4×10⁻⁶). Relative to exploratory sessions, exploitative sessions showed higher reward rate (β = 0.184, P = 0.0039). Consistent with the model, session-scale reward rate was higher during exploitative than exploratory sessions (58.4 ± 6.3% vs 49.9 ± 5.0%, P = 2.3×10^-7^; **Fig. 1G**). A paired comparison across rats confirmed this result (Wilcoxon signed-rank, P = 8×10^-8^; Fig. S3).

Session structure differed by strategy, as exploitative sessions had more trials (P = 1.6×10^-4^) and shorter trial durations (P = 9.4×10^-202^) (Fig. S3). Moreover, locomotion was slower under the exploratory strategy relative to exploitative sessions (P = 8.8×10⁻⁶; Fig. S3). Together, these results indicate a behavioural transition from outcome-reactive exploration early in training to an exploitation-biased strategy that tolerates isolated losses and yields higher reward later.

### Tonic cortico-cortical connectivity as a marker of task performance

To examine circuit dynamics during learning, we simultaneously recorded local field potentials from the prefrontal cortex (PFC), lateral entorhinal cortex (LEC), and dorsal CA1 (CA1d) of the hippocampus (**Fig. 2A**, Fig. S4). Power spectral density across the task revealed canonical frequency signatures for each region. CA1d exhibited prominent theta oscillations (4–10 Hz), consistent with known patterns during spatial navigation and active exploration (28, 37, 38). PFC showed dominant delta-band activity (2–4 Hz), a broad gamma shoulder (30–100 Hz), and a minor theta component. LEC also displayed strong delta power, a well-defined theta peak, and a narrower high-frequency shoulder in the fast-gamma (70–110 Hz) range (**Fig. 2B**).

**Figure 2.**
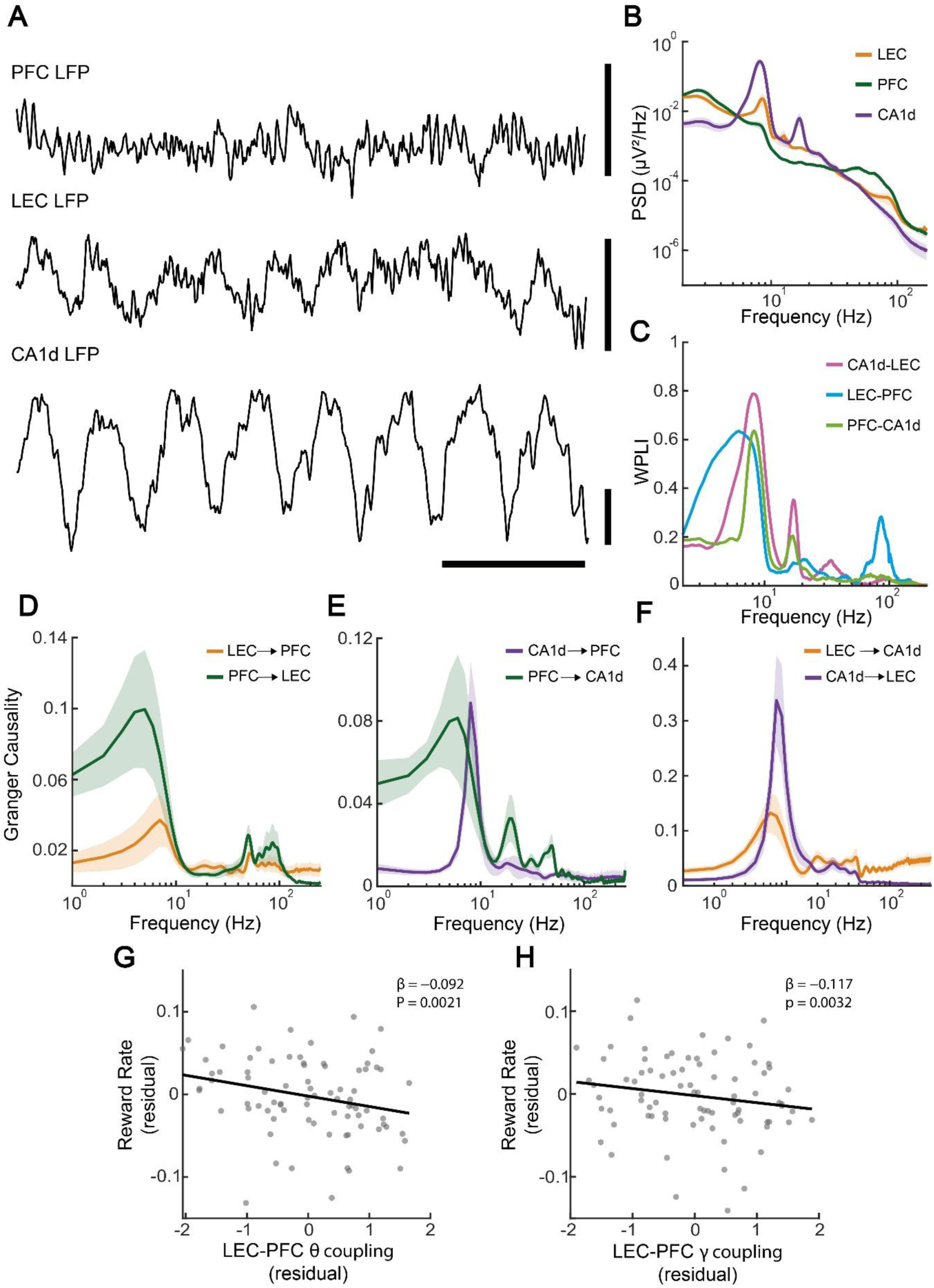
Field potentials and hippocampo–cortical coupling during task performance. **A,** Example broadband LFP recording from PFC, LEC, and CA1d. Scale bars: 500 µV (vertical), 250 ms (horizontal). **B,** Group-averaged normalized power spectra (PSD × frequency). For each animal, spectra were averaged across all included task sessions; the grand mean (± SEM across animals) is shown on log–log axes. **C,** Weighted phase–lag index (wPLI) spectra for each pair of regions. Curves are animal means; shaded bands are ± SEM across animals. Frequency axis is logarithmic. **D,** Spectral Granger causality (GC) by direction. For each animal, GC was averaged across sessions and then across animals (mean ± SEM; log-frequency axis). **E,** Session-wise residual reward rate (fraction of rewarded trials after controlling for session and strategy) versus residual LEC–PFC wPLI (after controlling for band-limited power and GC) in theta (θ, left) and fast-gamma (γ, right) bands (residualized and within-rat z-scored). Lines show partial fits from the full GLMM. Stronger LEC–PFC θ (β = −0.092, P = 0.0021) and fast-γ (β = −0.117, P = 0.0032) coupling were associated with lower reward rate across sessions; both effects survived FDR across neural predictors (q_FDR = 0.019).

Coordinated oscillations have been proposed as a mechanism for long-range communication between distributed neural circuits (28, 39). To test whether functional connectivity within the hippocampo-cortical circuit is structured by rhythmic activity, we used the weighted phase-lag index (wPLI) to quantify frequency-specific coupling between pairs of recorded regions (40, 41). As a first approach, we computed the spectra across the entire behavioral task, which revealed distinct global interaction profiles (**Fig. 2C**). The cortico-cortical pair (LEC–PFC) showed robust coupling in the delta–theta range (2–10 Hz), with an additional minor peak in the fast-gamma (70–110 Hz) band. CA1d–PFC connectivity was dominated by narrow-band theta coupling, with weaker coherence in the beta (10–20 Hz) range. Conversely, CA1d–LEC interactions spanned a broader theta range and uniquely expressed slow-gamma (20–50 Hz) coupling alongside beta synchrony. These patterns indicate that theta-range coordination is a shared motif across hippocampo-cortical circuits, while distinct high- and slow-gamma components differentiate specific pathways, thus suggesting that parallel frequency channels may support specialized forms of functional communication during learning (42, 43).

We next asked whether these rhythms supported directional interactions. Using frequency-resolved, non-parametric Granger causality computed per session and averaged across animals, we observed distinct asymmetries in the recorded regions. For the cortico-cortical pair (LEC–PFC), theta-band (peak at ∼6 Hz) influence was stronger in the PFC→LEC direction than LEC→PFC, with smaller components in the gamma band (**Fig. 2D**). For the hippocampo-cortical pairs, PFC→CA1d showed a similar theta peak (∼6 Hz) with weak gamma components, whereas the reverse CA1d→PFC direction expressed a sharper, faster theta peak (∼8 Hz) with negligible gamma contribution (**Fig. 2E**). The LEC–CA1d pair was dominated by a prominent CA1d→LEC theta drive (∼8 Hz), several-fold larger than the reverse, and with modest gamma coupling (**Fig. 2F**). These spectra align with the previous synchrony findings, highlighting the theta band as a common carrier of long-range coordination while revealing pathway-specific directionality. That is, theta oscillations dominated across pairs, whereas gamma effects were modest and pair-specific, thus suggesting theta and gamma frequencies as the principal channels of hippocampo-cortical communication during task performance.

To establish whether functional connectivity in this circuit relates to behavioral performance, we extended the linear mixed model by adding functional connectivity terms for the described LFP frequency bands and Granger causality as fixed effects (Table S3). Testing of fixed effects within the mixed model showed that only the cortico-cortical pair LEC–PFC was associated with reward rate, specifically theta (P = 0.019, FDR-controlled) and gamma (P = 0.019, FDR-controlled) frequencies. Neither hippocampo-cortical pairs (CA1d–LEC, CA1d–PFC) nor Granger causality reached significant effects (Table S3). To represent the relation between regional synchrony and task performance, we plotted the partial regression for the LEC–PFC wPLI in the theta (**Fig. 2G**) and gamma (**Fig. 2H**) bands and reward rate. In both oscillatory bands the slope was negative, indicating that, controlling for training and strategy, sessions with stronger LEC–PFC synchrony exhibited lower reward rate, consistent with the linear model estimates. Because these metrics were estimated across the entire session, the effect likely reflects tonic cortico-cortical coupling. Taken together, these data point to cortico-cortical (LEC–PFC) theta-gamma coupling as a consistent negative marker of behavioural performance.

### Hippocampo-cortical theta oscillations gate cortical gamma bursts and bias spike timing

Although hippocampo–cortical connectivity was not directly predictive of reward rate at the session scale (Table S3), Granger causality nonetheless indicated a pronounced hippocampal drive to cortex in the theta band, motivating a mechanistic analysis. We therefore asked whether hippocampal theta waves can structure cortical gamma bursts (**Fig. 3A**). Cross-region phase–amplitude coupling (PAC) revealed that CA1d theta phase significantly modulated gamma amplitude in both LEC and PFC, yielding robust PAC maps (**Fig. 3B**) and imprinting significant phase modulation for cortical gamma bursts, including LEC fast-gamma (Rayleigh test, P = 0.02; **Fig. 3C**), PFC fast-gamma (Rayleigh test, P = 6.5×10^-5^; **Fig. 3D**) and LEC slow-gamma (Rayleigh test, P = 2.6×10^-4^; **Fig. 3E**).

**Figure 3.**
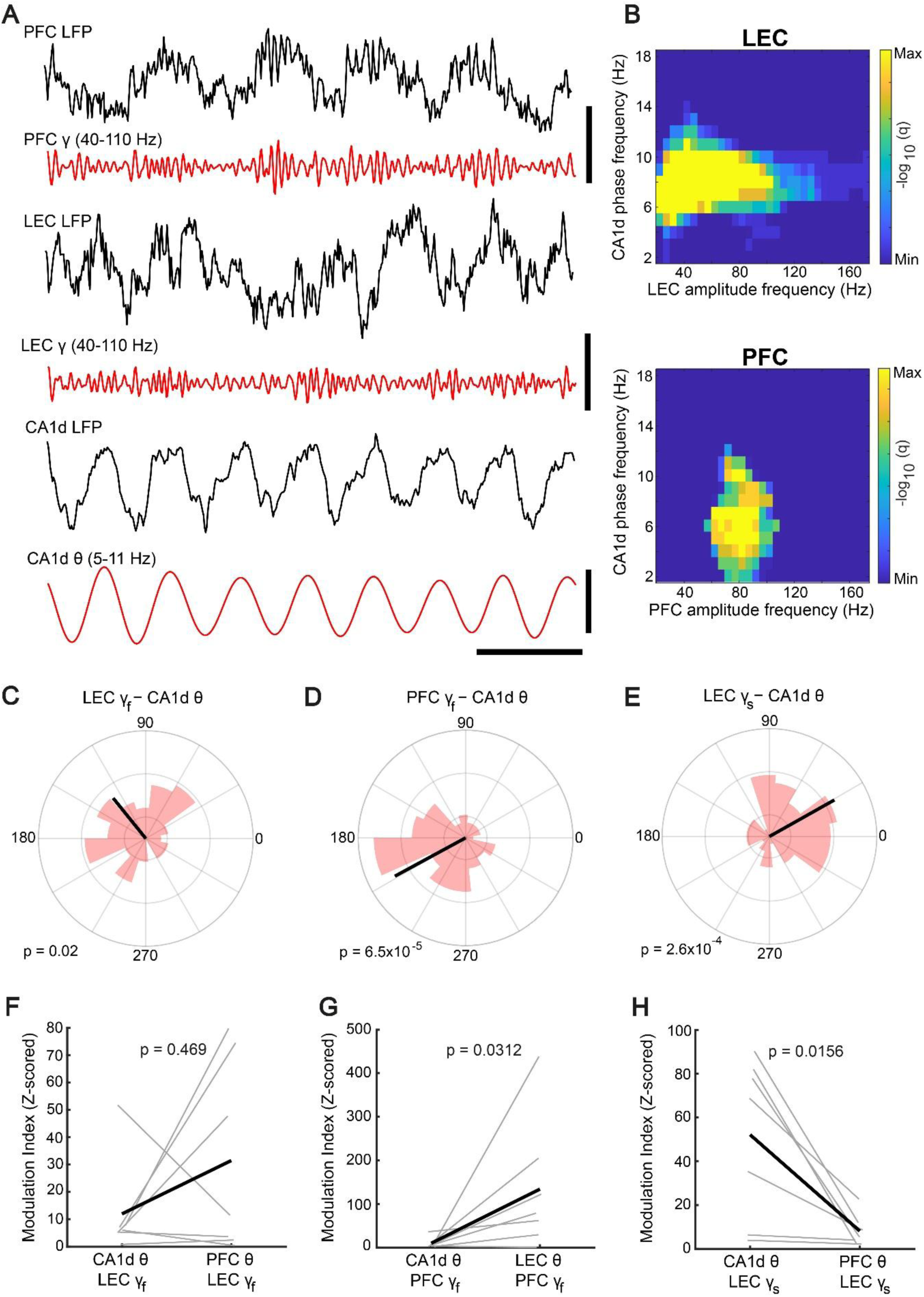
Theta–gamma interactions in the hippocampo-cortical circuit. **A,** Simultaneous LFP recordings from PFC, LEC, and dorsal CA1 (black). For PFC and LEC, the γ-band filtered signal (40–110 Hz) is overlaid in red; for CA1d the red trace is θ (5–11 Hz). Scale bars: 300 µV (PFC/LEC), 1 mV (CA1d), 200 ms (all). **B,** Group phase–amplitude coupling (PAC) significance maps for LEC (top) and PFC (bottom). Each panel shows the phase-frequency (CA1d θ on the y-axis) by amplitude-frequency grid, expressed as −log10(q); q-values are Benjamini–Hochberg FDR–corrected Wilcoxon signed-rank tests across animals. **C–D,** PAC (modulation index, z-scored relative to within-animal null) for fast-γ amplitude (70–110 Hz) in LEC (C) or PFC (D), driven by θ from two sources (CA1d vs LEC/PFC). **E,** PAC for slow-γ amplitude in LEC (20–50 Hz) driven by θ from CA1d vs PFC (black line represents mean over animals). PAC was stronger for one source (P = 0.0156, Wilcoxon signed-rank). Points are animals; lines pair within-animal comparisons; black symbols are mean ± SEM. In LEC (C) PAC was not different for CA1d-θ vs PFC-θ drive (P = 0.469, Wilcoxon signed-rank). In PFC (D) PAC depended on θ source (P = 0.0312). **F–H,** Circular histograms of the preferred phase of γ bursts recorded in one region relative to θ band in another region. Phase histogram of preferred phases across bursts (pink) and population mean vector (black), whose length corresponds to the population phase alignment (|U|, defined as the magnitude of the mean of unit-normalized phase resultant vectors). F, LEC; |U| = 0.2414, Rayleigh P = 0.0196, z = 3.9037. G, PFC; |U| = 0.374, Rayleigh P = 6.47×10⁻⁵, z = 9.3711). H, LEC; |U| = 0.3467; Rayleigh P = 2.63×10⁻⁴; z = 8.05.

We next compared candidate theta sources for cortical gamma modulation. In LEC, PAC strength did not differ between CA1d-theta→LEC-fast-gamma and PFC-theta→LEC-fast-gamma (P = 0.469; **Fig. 3F**), suggesting that both hippocampal and cortical theta provide comparable temporal reference signals for LEC fast-gamma. In contrast, in PFC, CA1d-theta→PFC-fast-gamma though significant, was weaker than LEC-theta→PFC-fast-gamma (P = 0.0312; **Fig. 3G**), consistent with a stronger cortico–cortical influence on PFC gamma timing. Finally, hippocampal theta also organized slow-gamma activity in LEC, with stronger CA1d-theta→LEC-slow-gamma coupling than PFC-theta→LEC-slow-gamma coupling (P = 0.0156; **Fig. 3H**). Together, these results indicate that CA1d theta provides a reliable temporal scaffold for cortical gamma bursts, with hippocampal influence most prominent for LEC slow-gamma and cortico–cortical influence more prominent for PFC fast-gamma.

Hippocampo-cortical modulation was reliable over time, as it did not change across training and strategy (Table S4). Directional influences, however, were influenced by training, particularly in the hippocampo-entorhinal pathway. Indeed, the CA1d→LEC theta drive increased with training (β = 0.136, P = 0.019) while the reverse direction remained stable (P = 0.278). In slow-gamma, Granger causality increased bidirectionally (CA1d→LEC, β = 0.119, P = 0.014; LEC→CA1d, β = 0.107, P = 0.002). Together, these findings indicate that hippocampal theta waves enable temporal windows that bias cortical gamma bursts, consistent with a regulatory role over cortico-cortical dynamics. Importantly, adding PAC to the reward rate model did not improve association beyond training and connectivity (Table S5), implying that inter-regional theta–gamma coordination is mechanistic but not a session-scale marker of task performance. Thus, theta-gated gamma coordination appears to coexist with cortico-cortical coupling, but it does not add predictive value for reward rate once training and connectivity are controlled.

To test whether theta rhythms can also organize spike timing across regions, we aligned cortical spikes to the instantaneous phase of band-limited field potentials and quantified the resulting phase distributions. LEC theta phase biased PFC spike timing (Rayleigh test, P = 0.033; **Fig. 4A**), and CA1d theta phase also significantly modulated PFC spiking (Rayleigh test, P=5.83×10^-12^; **Fig. 4B**). To compare the unit phase-locking strength between regions, we quantified the median unit mean resultant length (MRL) within each session. We found that CA1d-driven modulation of PFC units, though significant, was weaker than LEC-driven modulation (P = 0.00261, **Fig. 4C**). In the reverse direction, PFC theta phase significantly modulated LEC unit discharge (Rayleigh test, P = 2.4×10^-9^; **Fig. 4D**), and CA1d theta phase also biased LEC spiking (Rayleigh test, P = 1.2×10^-5^; **Fig. 4E**). In this case, the CA1d-driven modulation of LEC units was stronger than PFC-driven modulation (P = 0.00458, **Fig. 4F**). Finally, the preferred CA1d theta phase at which gamma burst and spike timing were modulated was comparable within each hippocampo–cortical pathway, consistent with overlapping temporal windows of interaction (Watson-Williams test, CA1d-theta→LEC-fast-gamma vs CA1d-theta→LEC-units, P = 0.839; CA1d-theta→PFC-fast-gamma vs CA1d-theta→PFC-units, P = 0.242).

**Figure 4.**
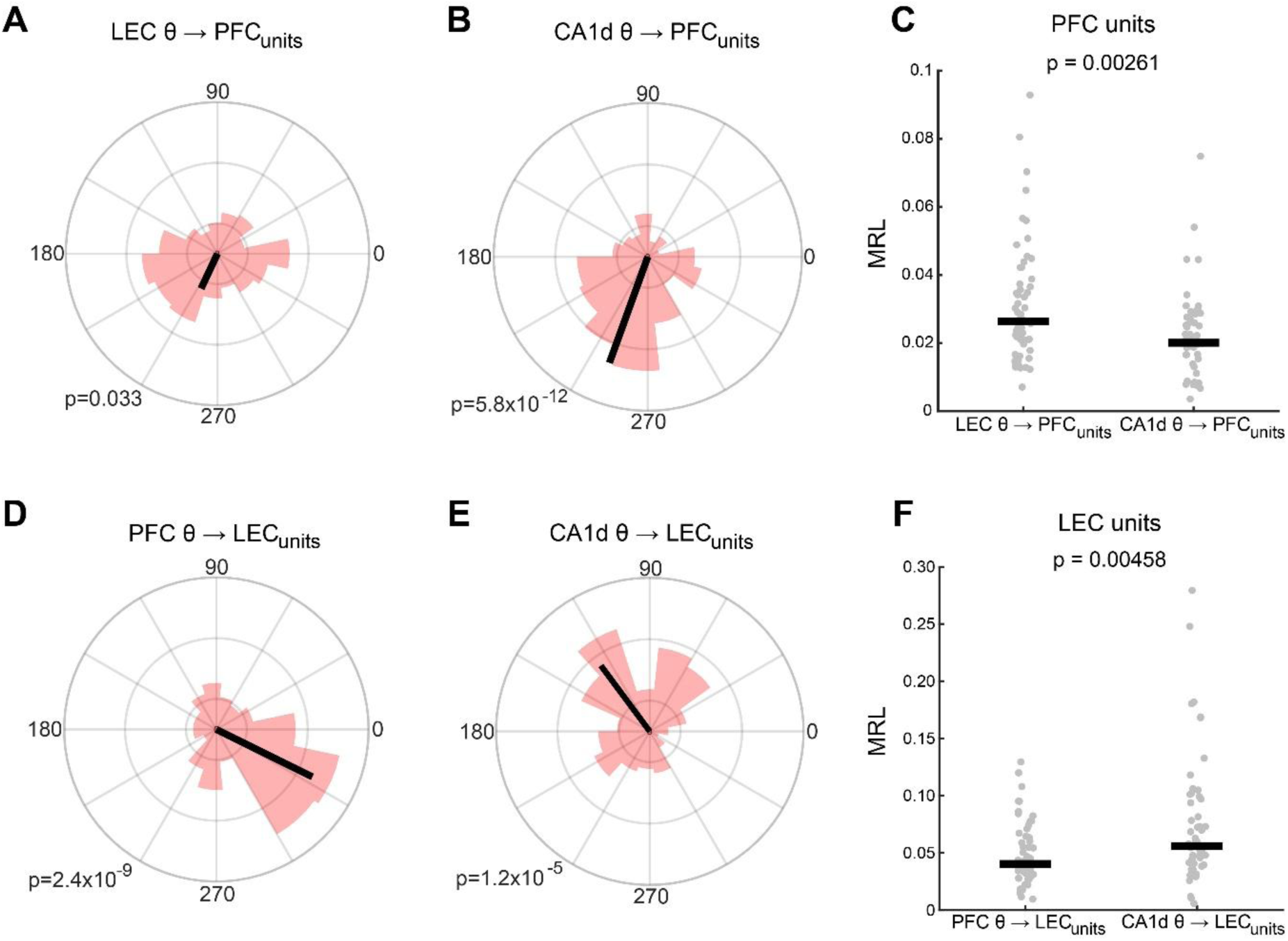
Cross-regional theta modulation of spike timing. Rose plots show the preferred phase of spikes recorded in one region relative to band-limited LFP in another region. Phase histogram of preferred phases across units (pink) and population mean vector (black), whose length corresponds to the population phase alignment (|U|, defined as the magnitude of the mean of unit-normalized phase resultant vectors). **A**, PFC spikes relative to LEC θ (|U| = 0.127; Rayleigh test, P = 0.033, z = 3.40). **B**, PFC spikes relative to CA1d θ (|U| = 0.366; Rayleigh test, P = 5.83 × 10⁻¹², z = 25.04). **D**, LEC spikes relative to PFC θ (|U| = 0.352; Rayleigh test, P = 2.40 × 10⁻⁹, z = 19.28). **E**, LEC spikes relative to CA1d θ (|U| = 0.267; Rayleigh test, P = 1.19 × 10⁻⁵, z = 11.17). Session-level comparisons of unit phase-locking strength, quantified as the median unit mean resultant length (MRL) within each session. Each dot represents one session; black bars indicate the median across sessions. **C**, PFC unit locking to LEC θ versus CA1d θ (Mann–Whitney U test, P = 0.00261). **F**, LEC unit locking to PFC θ versus CA1d θ (Mann–Whitney U test, P = 0.00458).

In addition, in the entorhinal–hippocampal pathway, Granger causality revealed bidirectional slow-gamma influence, which was paralleled by reciprocal phase-locked spiking in the same band (CA1d-slow-gamma→LEC units, P = 6.6×10^-5^; LEC-slow-gamma→CA1d-units, P = 8.9×10^-7^). Together, these findings indicate that hippocampal theta waves provide a common temporal reference frame that gates cortical gamma bursts and aligns spike timing across the cortex. Theta–gamma coordination and phase-locked unit discharge are therefore robust circuit mechanisms for structuring information flow, even though they do not themselves correlate with session-scale performance.

### Peri-goal neuronal spiking tracks locomotor dynamics and segregate into prefrontal ramping and hippocampo–entorhinal suppression

Neuronal firing rates were affected by training, specifically, neuronal spiking in the PFC decreased with training (P = 0.027, **Fig. 5A**), while CA1d (P = 0.329, **Fig. 5B**) and LEC (P = 0.103, **Fig. 5C**) unit activity remained consistent over time. To connect cortical oscillations to neuronal output at specific behavioral events, we aligned normalized peri-goal firing rates to the nose-poke in the reward port and classified units as ‘activated’ (peak increase) or ‘inhibited’ (peak decrease) (Fig. S5). Latency-sorted heatmaps revealed robust, region-specific modulation. In PFC, modulation was dominated by activation (134/232 units, 57.8%), forming a population ramp that began briefly before the nose-poke and remained tonically elevated afterward. Average traces showed a stable separation between activated and inhibited populations during the goal-reaching interval, consistent with sustained engagement of PFC neurons around choice execution (**Fig. 5D**). In LEC the balance reversed (56/184 activated, 30.4%), with a small population showing brief activation during goal-approach and a larger fraction inhibiting upon goal-reaching, followed by a slow recovery (**Fig. 5E**). CA1d showed the strongest suppression (51/171 activated, 29.8%), with a sharp trough centered on the nose-poke and a gradual recovery to baseline thereafter (**Fig. 5F**). Direct comparison of population spiking confirmed that CA1d exhibited a stronger suppression upon goal-reaching, whereas LEC and PFC remained closer to baseline levels (Fig. S6). Across regions, the latency-sorted discharge profiles indicate a consistent temporal sequence, in which a cortical build-up led by PFC precedes the nose-poke, a briefer overlapping suppression in LEC, and a pronounced hippocampal suppression tightly locked to goal-reaching.

**Figure 5.**
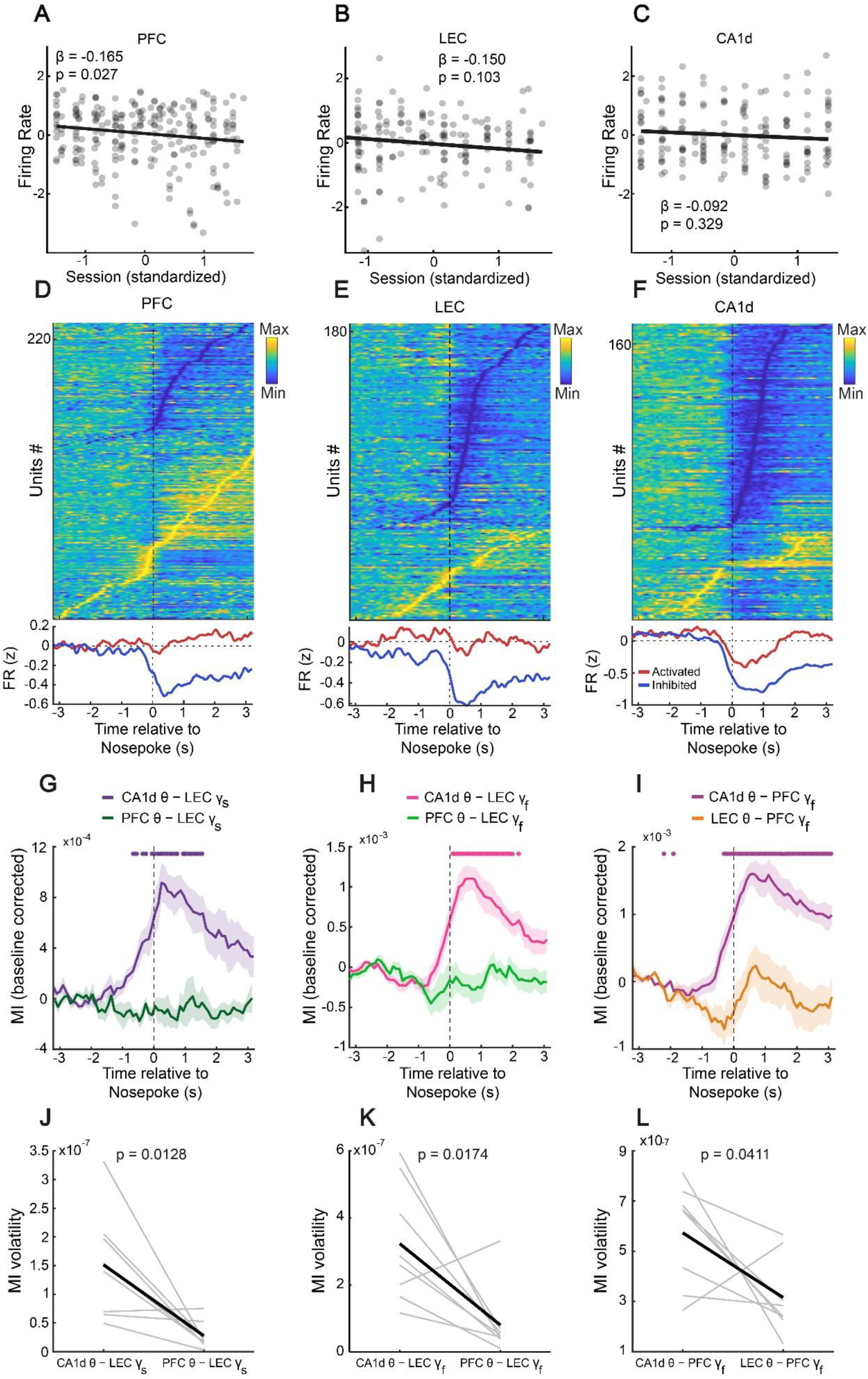
Peri-goal firing patterns and time-resolved theta–gamma coupling in the hippocampo-cortical cirtcuit. A-C, Unit firing rates (log-transformed, then z-scored within unit) versus standardized training session. Points depict individual units; lines show fixed-effect fits from a linear mixed model (random intercepts for rat and unit; covariates included strategy and session-level unit counts). A: PFC firing decreased with training (β = −0.165, P = 0.027); B-C: LEC and CA1d showed no significant trends (LEC, β = −0.15, P = 0.103; CA1d, β = −0.092, P = 0.329). **D–F,** Peri-event heat maps of unit activity in PFC (D), LEC (E), and CA1d (F), aligned to nosepoke onset (t = 0). Rows are units pooled across rats and sessions, z-scored within unit, and sorted by the latency of maximal modulation. Activated (red) and Inhibited (blue) neurons are defined by each unit’s peak positive vs negative deviation. Bottom traces show the mean ± SEM per group. PFC (activated, n = 134; inhibited, n = 98), LEC (activated, n = 56; inhibited, n = 128), and CA1d (activated, n = 51; inhibited, n = 120). **G–I,** Time-resolved phase–amplitude coupling (PAC) around nosepoke. Curves are the baseline-corrected modulation index (MI), plotted as mean ± SEM across animals. G: LEC slow-γ amplitude coupled to CA1d θ vs PFC θ. H: LEC fast-γ amplitude coupled to CA1d θ vs PFC θ. I: PFC fast-γ amplitude coupled to CA1d θ vs LEC θ. Asterisks mark time bins where baseline-corrected MI differs from zero (one-sample t-test across animals, Benjamini–Hochberg FDR, q = 0.05). **J-L,** Volatility of PAC dynamics for the same pairs (G–I), defined per animal as the temporal variance of baseline-corrected MI in the peri-nosepoke window. Paired two-sided t-tests compare configurations (gray lines, individual animals; black lines, population average): J, LEC slow-γ (CA1dθ→LECγ-slow vs PFCθ→LECγ-slow, P = 0.0128); K, LEC fast-γ (CA1dθ→LECγ-fast vs PFCθ→LECγ-fast, P = 0.0174); and L, PFC fast-γ (CA1dθ→PFCγ-fast vs LECθ→PFCγ-fast, P = 0.0411).

Cross-regional theta–gamma coupling has been proposed as a routing mechanism in cortical networks (28, 44). We therefore studied peri-goal hippocampal theta interacts with cortical gamma bursts, when trial information must be integrated and updated. Phase–amplitude coupling revealed a noticeable increase in hippocampo–cortical coordination upon goal-reaching, whereas cortico–cortical coupling remained comparatively stable. Indeed, CA1d theta phase dynamically gated slow-gamma bursts in LEC, with coupling rising during goal-approach, peaking shortly after goal-reaching, and then slowly decaying toward baseline, while PFC-theta→LEC-fast-gamma coupling remained stable over the same interval (**Fig. 5G**). Likewise, CA1d theta phase tightly controlled fast-gamma bursts in both LEC (**Fig. 5H**) and PFC (**Fig. 5I**), following a similar temporal profile to the CA1d-theta→LEC-slow-gamma interaction. Moreover, we quantified these dynamics at the single-animal level. For each rat, we computed the variance of theta–gamma MI across the peri-goal interval, providing an index of how strongly coupling fluctuated over time rather than its average strength. In every case, cross-frequency coupling was more volatile for hippocampo-cortical pathways than for cortico-cortical connections (**Fig. 5J–L**). This pattern indicates that hippocampal theta oscillations transiently organize cortical gamma activity around goal-reaching, even though this time-locked gating does not itself improve performance association at the broader session-scale.

To assess whether the peri-goal firing patterns reflected locomotor dynamics, we correlated the time course of population spiking with running speed. For each recorded region we normalized the firing rates of activated and inhibited units and computed the Pearson correlation between them and the mean speed profile during the peri-goal interval. In PFC, activated units showed a robust negative correlation with speed (r = −0.48, P = 2×10^-4^), whereas inhibited units were strongly positively correlated (r = 0.87, P = 2×10^-4^) (**Fig. 6A**). In LEC, both populations also covaried with speed, but the relation was stronger for inhibited units (activated: r = 0.35, P = 2×10^-4^; inhibited: r = 0.82, P = 2×10^-4^) (**Fig. 6B**). A similar pattern was observed in CA1d, where the inhibited units tracked speed closely (r = 0.84, P = 2×10^-4^), while the activated population showed only a weak correlation (r = 0.14, P = 0.009) (**Fig. 6C**). Thus, peri-goal suppression in LEC and CA1d largely mirrors the deceleration profile, whereas the prefrontal activated population increases as animals slow down, indicating that its ramping is not a simple reflection of speed but is instead aligned to the goal-approach and goal-reaching.

**Figure 6.**
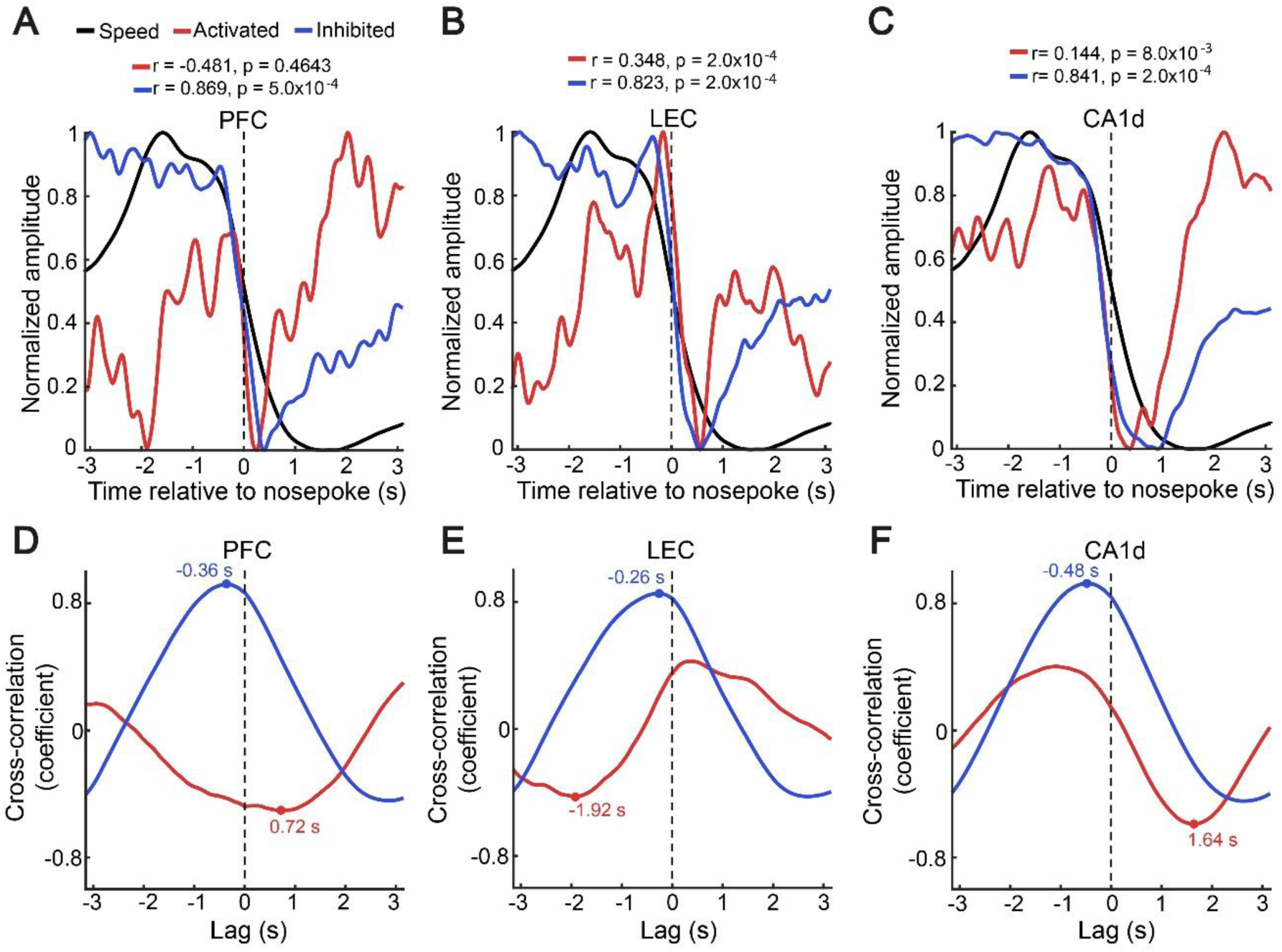
Peri-goal population spike-timing covaries with locomotor speed. A-C, Cross-correlation between mean peri-nosepoke speed and population-averaged z-scored firing rates for the activated (red) and inhibited (blue) groups in each recorded area. Curves show coefficients as a function of lag (s); negative lags mean neural activity precedes speed. Dots mark the lag of the peak absolute correlation. A, PFC: activated, max. coeff. = 0.503 at +0.72 s (P = 0.4578); inhibited, max. coeff. = 0.922 at −0.36 s (P = 0.0005). B, LEC: activated, max. coeff. = 0.427 at −1.92 s (P = 1); inhibited, max. coeff. = 0.853 at −0.26 s (P = 0.0005). C, CA1d: activated, max. coeff. = 0.590 at +1.64 s (P = 0.6687); inhibited, max. coeff. = 0.926 at −0.48 s (P = 0.0005). **D-F,** Normalized peri-nosepoke time courses for speed (black) and population activity (red, activated; blue, inhibited). The scalar Pearson correlation showed that inhibited populations were consistently more correlated with speed that activated populations over the peri-nosepoke window (permutation test via time shuffling). D, PFC: activated, r = −0.481, P = 0.0002; inhibited, r = 0.869, P = 0.0002. E, LEC: activated, r = 0.348, P = 0.0002; inhibited, r = 0.823, P = 0.0002. F, CA1d: activated, r = 0.144, P = 0.01; inhibited r = 0.841, P = 0.0002.

Given the strong correlation of inhibited units across regions with movement deceleration, we then asked whether neuronal changes followed speed or instead anticipated it. Using cross-correlation between population spiking and velocity, combined with a surrogate test on the maximal absolute correlation, we found that in all three regions the inhibited populations preceded changes in speed. In PFC, the strongest correlation occurred when neuronal activity led speed by about 0.36 s (maximal coefficient = 0.92, P = 5×10^-5^), indicating that decreases in discharge activity in the inhibited ensemble reliably occurred before subsequent slowing (**Fig. 6D**). A similar anticipatory pattern was observed in LEC (maximal coefficient = 0.85, P = 5×10^-5^, **Fig. 6E**) and CA1d (maximal coefficient = 0.93, P = 5×10^-4^, **Fig. 6F**), where the inhibited populations led velocity by roughly 0.26 s and 0.48 s, respectively, whereas correlations for the activated populations were weaker and not significant. Together with the peri-event profiles, these analyses show that hippocampal and entorhinal suppression is tightly coupled, and slightly ahead of, locomotor deceleration, while PFC maintains an additional sustained spiking component that remains elevated beyond what can be accounted for by changes in speed.

Taken together, these results indicate that goal-reaching is accompanied by a stereotyped reconfiguration of the circuit, so that a prefrontal population ramps and remains engaged after goal-reaching, while hippocampal and entorhinal populations enter a brief, speed-linked suppression that slightly precedes deceleration during goal-approach. This peri-goal state thus reflects a shift toward prefrontal dominance, embedded within a hippocampo–entorhinal channel that aligns cortical dynamics to the timing of behavior.

## Discussion

We have studied how a hippocampo–cortical circuit linking CA1d, LEC and PFC supports learning under uncertainty in a probabilistic reversal task. Behaviorally, training drove a gradual shift from outcome–reactive exploration toward an exploitation–biased strategy that tolerated isolated losses and produced higher reward rates. Taking advantage of simultaneous recordings across the three regions, we then used mixed-effects models to identify a neural marker of task performance at the session-scale and dissociate it from others tied to specific behavioral events. Three main conclusions emerge in this study. First, tonic cortico–cortical synchrony in the theta and fast–gamma bands is a marker of task performance (45, 46), as stronger LEC–PFC coupling consistently associates with lower reward rate after controlling for training and strategy. Second, hippocampal theta acts as a directional driver that gates cortical gamma bursts and spike timing (47, 48), particularly in the hippocampo-entorhinal pathway. Third, during the peri-goal interval the circuit displays a state in which prefrontal activity ramps (49) while hippocampal and entorhinal populations are suppressed (50), with peri–goal dynamics closely tracking locomotor deceleration.

One significant result is the negative relation between tonic theta-gamma LEC–PFC synchrony and session-scale reward rate (51). When the principal frequency bands and pairwise connections were included in the behavioral model, only theta and fast–gamma connectivity in the cortico–cortical pair survived as significant effects, and higher coupling was consistently associated with poorer performance. This effect persisted after accounting for training and strategy, and was absent in hippocampo–cortical connections and in Granger directionality. These findings argue against the simple idea that synchrony directly improves performance and instead suggest the opposite (at least, for this probabilistic reversal learning task), so that strong LEC–PFC coupling reflects a cortical state that is maladaptive for rapid adjustment to changing contingencies (47, 51). One possibility is that excessive coherence reflects over–stabilization of a current decision strategy or context representation, making it harder to reassign credit when the latent state changes (52, 53). Here, we define a marker as a metric that predicts task performance while controlling for training and strategy. In this context, tonic LEC–PFC synchrony operates as a marker of reward rate, rather than as a direct mechanism driving performance improvement.

In contrast, the hippocampal theta rhythm exerted a clear directional influence on entorhinal and prefrontal dynamics that was strongly modulated by learning (54–56). Granger analysis revealed a prominent CA1d→LEC theta drive that increased with training, while the reverse LEC→CA1d influence remained invariant. Slow–gamma interactions were specific for this pathway and became bidirectional with learning, consistent with a progressive strengthening of hippocampo–entorhinal dialogue. Cross-frequency coupling further showed that CA1d theta phase robustly gated both slow– and fast-gamma bursts in LEC and modulated fast–gamma bursts in PFC, and these effects were stable across strategies. Together, these results support a model in which hippocampal theta provides temporally precise windows for entorhinal and prefrontal processing (30, 48, 57), particularly as animals become more familiar with the task structure. Importantly, however, adding PAC metrics to the reward rate model did not improve association beyond LEC–PFC connectivity and training. Thus, theta–gamma coordination appears as a mechanistic scaffold for organizing cortical computations rather than as an additional session–scale marker of task performance.

Our results show a dissociation between the cross-cortical coupling marker of reward rate and the hippocampal theta mechanism for temporal organization. Also, the entorhinal pathway plays a special role in this framework. LEC sits at the interface between hippocampus and PFC, encoding local option identity and recent outcomes (20) while funneling information into CA1d (58). Our data suggest that LEC participates in both sides of the dissociation. On one hand, its long–range coupling with PFC provides the tonic synchrony that tracks performance (59). On the other hand, LEC gamma bursts are tightly gated by CA1d theta and display strong theta–gamma coupling (57, 60), especially in the slow–gamma band where Granger influence is bidirectional. This position in the circuit is ideally suited for LEC to act as a conditional gate, so that when strongly synchronized with PFC it may stabilize a current belief about the environment, whereas when driven by CA1d theta it may be more engaged in updating value estimates or contextual representations.

Also, our peri–goal analyses further clarified how these mechanisms operate at the lower level of neuronal spiking populations. Aligning unit activity to the nose–poke revealed a clear segregation of dynamics across the circuit. In PFC, the majority of units classified as activated showed a sustained ramp that began shortly before goal–reaching and remained elevated afterward, while inhibited units decreased firing in parallel with locomotor slowing. Several studies have shown ramping activity in PFC during behaviourally relevant moments, like choice-selection or target-approaching (55, 61–64). In LEC and CA1d, inhibition dominated, as both regions exhibited a sharp suppression of activity during peri-goal intervals, followed by a slow recovery, consistent with a transient reset of hippocampo–entorhinal activity during outcome evaluation (65, 66). These neural activity patterns are consistent with a model in which prefrontal populations remain engaged in representing the chosen option and outcome evaluation, while hippocampal and entorhinal ensembles briefly disengage and likely reset.

This interpretation is further consolidated by quantifying the relationship between peri–goal neuronal spiking and locomotor speed. Across all three regions, inhibited populations tightly tracked the deceleration profile, whereas the activated PFC ensemble was anti–correlated with the animal speed. Crucially, cross–correlation analyses showed that the peak correlations for the inhibited populations occurred when neuronal spiking led changes in speed by several hundred milliseconds. These anticipatory relationships argue that the peri–goal suppression is unlikely to be a passive reflection of speed slowing, but instead contributes to the control of movement and state transitions. At the same time, the sustained PFC spiking ramp persists beyond what can be accounted for by speed, indicating that prefrontal neurons are likely to encode goal-approach (61, 67) and outcome-assessment (14, 68), that are partially decoupled from locomotor kinematics. Together with the oscillatory data, these findings support a model in which hippocampo-entorhinal ensembles participate in shaping upcoming deceleration and state change, while PFC maintains a higher–order representation of choice and expected outcome (69).

These results also have implications for theoretical accounts of credit assignment in recurrent circuits. For example, reinforcement learning models with latent states require that outcome information be routed back to the representations that generated the choice (70–72). The tonic LEC–PFC coupling that associates with performance may reflect the current credit assignment configuration of the cortical network, with strong synchrony corresponding to a more rigid mapping between states and actions. In parallel, hippocampal theta–gamma coordination, especially in the CA1d–LEC pathway, may provide temporal windows that bind together entorhinal content and hippocampal state representations, enabling efficient updating when contingencies change. The peri–goal suppression and anticipatory coupling to deceleration suggest that these updates are embedded within finely-tuned transitions in motor and cognitive states. Further experiments will have to be developed to test this model, which currently fall beyond the scope of the current study.

Several limitations of the present work should be acknowledged. First, our analyses are correlational and rely on local field potentials and multiunit spiking. Demonstrating causality will require targeted manipulations of specific frequency bands or anatomical pathways, for example closed–loop perturbations of CA1d theta or LEC–PFC synchrony. Second, our emphasis is necessarily at the mesoscale, given that our recording yield is pretty modest compared with modern high-density probes (73, 74), so we sample tens rather than thousands of neurons. This constrains seriously our ability to resolve fine-grained cell-type or population codes. Finally, our models purposely focus on session-averaged reward rate and decision strategy; relating the observed rhythms to trial-by-trial value estimates or belief updates remains an important goal for future work. Despite these caveats, the present findings advance our understanding of how hippocampus, entorhinal cortex and prefrontal cortex jointly support learning under uncertainty. By separating a cortical synchrony marker of performance from a hippocampal theta mechanism that organizes cortical gamma bursts and spike timing, and by showing how these processes unfold around goal–reaching and locomotor transitions, we provide a circuit–level framework for how latent state inference, strategy adaptation and motor control are integrated within a hippocampo–cortical circuit.

## Methods

### Animals

Nine adult male Sprague–Dawley rats (postnatal day 40–60; 300–350 g) were obtained from the Center for Innovation on Biomedical Experimental Models (CIBEM; https://cibem.bio.puc.cl/) at the Pontificia Universidad Católica de Chile. Animals were housed in a temperature-controlled room (22 ± 1 °C) on a 12 h light/dark cycle (lights on 07:00) with food and water ad libitum. All procedures were approved by the Scientific Ethical Committee for the Care of Animals and the Environment (CEC-CAA; protocol 240701014). All methods complied with relevant guidelines and regulations and are reported in accordance with the ARRIVE guidelines (https://arriveguidelines.org).

### Habituation

Animals were first acclimated to the housing room for three days, then handled by the experimenter daily for an additional 3–5 days. They were subsequently habituated to the recording environment by being placed on the task maze in the recording room for 30–60 min per day for about one week.

### Behavioral task

Rats performed a self-paced two-armed bandit in a maze with a start box, a central stem, and two lateral goal ports (nose-pokes). On each trial animals exited the start box, ran along the stem, and chose one of the two ports to trigger potential reward delivery; they then returned to the start box to initiate the next trial. The task was organized in blocks in which the reward rate associated with each option was fixed, and block identities switched without explicit cues. Outcomes were delivered stochastically on every trial, so animals could only infer block transitions by trial-and-error. Training comprised single daily sessions; across days, the number of trials per session and average running speed were approximately stable. Position tracking provided linear speed and distance measures used both to describe behavior and to define analysis epochs along the trajectory (start-box, stem, approach, nose-poke, return).

### Recording Implant Assembly

The recording microdrive was designed in Autodesk Fusion and 3D-printed on an F410 printer. A custom microdrive carried 16 or 32 tungsten, standard tapered-tip electrodes (MicroProbes, USA; 5 MΩ), positioned to target the regions of interest according to a rat stereotaxic atlas. Each electrode was wired to an EIB-36 PCB (36 channels, including ground and reference). Ground and reference leads were connected to stainless-steel skull screws placed during surgery, and the entire assembly was enclosed in copper mesh to minimize electrical noise.

### Stereotaxic Surgery

Rats were anesthetized with isoflurane (4% for induction, 1.5–2% for maintenance) and secured in a stereotaxic frame (Stoelting). Body temperature was held at 35–37 °C with a homeothermic blanket, and animals received hourly glucosaline (0.9% NaCl, 2.5% dextrose). After a scalp incision, three ∼1 mm craniotomies were made in the right hemisphere at predetermined coordinates (all in mm): CA1d, −4.0 AP, 2.0 ML, 2.4 DV; LEC, −6.8 AP, 6.3 ML, 7.4 DV; PFC, 2.5 AP, 0.4 ML, 4.2 DV. Two additional craniotomies anterior to bregma accommodated ground and reference electrodes, and four more (two contralateral parietal, one ipsilateral parietal, one posterior to lambda) secured the implant with screws. The dura was removed at the target sites, cortical surfaces were kept moist with mineral oil, and electrodes were inserted carefully. Craniotomies were sealed with silicone elastomer or wax, and the microdrive was fixed with dental acrylic. Postoperative care consisted of daily subcutaneous enrofloxacin (5 mg/kg) and meloxicam (2 mg/kg) for three days. Animals recovered for at least seven days before recordings.

### Electrophysiological Recordings

After recovery, electrophysiological recordings were performed for 10 consecutive days during the light phase inside a Faraday-shielded enclosure. Rats were placed on the task maze for roughly 60–min sessions. The EIB-36 board was connected via a 16- or 32-channel headstage (Intan Technologies) to an Intan RHD amplifier, with one electrode used as the reference. Video, synchronized to the amplifier clock, assisted brain-state identification. Signals were sampled at 20 kHz with RHX Data Acquisition (Intan Technologies) and converted to MATLAB using the LAN toolbox for analysis.

### Weighted phase-lag index (wPLI)

Phase synchronization between pairs of LFP signals was quantified using the debiased weighted phase-lag index (wPLI), a measure of consistent non-zero phase-lag coupling that reduces spurious synchrony arising from volume conduction and common reference effects (39). Continuous LFP signals were segmented into non-overlapping temporal windows of fixed duration. Spectral estimates were computed for each window using the FieldTrip toolbox, and phase synchrony was quantified from the resulting cross-spectral density. Connectivity estimates were obtained using the debiased formulation of wPLI, which corrects for sample-size–dependent bias.

### Granger causality

Directional interactions between recorded brain regions during maze sessions were quantified using frequency-resolved Granger causality (GC) (41). Each recorded region were normalized to place channels on a comparable scale. For each unique pair of recorded areas, signals were locally detrended to reduce slow fluctuations before causal analysis. Granger causality was estimated in the frequency domain, yielding directed GC spectra for both directions of each area pair.

### Cross-frequency phase–amplitude coupling

Cross-frequency interactions were quantified using phase–amplitude coupling (PAC), assessed via comodulograms. Analyses were performed on LFP signals. Temporal padding was applied to minimize edge-related artifacts. Signals were locally detrended prior to filtering. Low-frequency phase signals were obtained by band-pass filtering the LFP into overlapping phase-frequency bands spanning 1–17 Hz (2-Hz bandwidth, 1-Hz steps), followed by extraction of instantaneous phase via the Hilbert transform. High-frequency amplitude signals were obtained by band-pass filtering the same LFP into overlapping amplitude-frequency bands spanning 20–170 Hz (4-Hz bandwidth, 5-Hz steps), followed by extraction of the instantaneous amplitude envelope. θ bands followed regional definitions (PFC/LEC: 6–10 Hz; CA1d: 4–11 Hz). Phase–amplitude coupling was quantified using the modulation index (MI), computed by comparing the empirical distribution of high-frequency amplitude across phase bins to a uniform distribution. Phase was discretized into 18 equally spaced bins spanning −π to π. For each phase–amplitude frequency pair, the MI was computed, yielding a two-dimensional comodulogram describing coupling strength across frequency pairs. Statistical significance of PAC was assessed using a surrogate-based null distribution constructed independently for each session. Surrogate comodulograms were generated using a circular time-shifting procedure that preserves the spectral and temporal structure of the phase and amplitude signals while disrupting their temporal alignment. Specifically, the high-frequency amplitude envelope was circularly shifted by a random temporal offset relative to the low-frequency phase signal, while the phase time series was left unchanged. To avoid trivial correlations arising from short temporal shifts, the minimum allowed shift for each amplitude-frequency band was determined from the temporal autocorrelation structure of the amplitude envelope. The minimum shift was defined as the first lag at which the autocorrelation dropped below 1/e, indicating approximate temporal decorrelation. 100 surrogate realizations were generated per session to form a session-specific null distribution of comodulograms.

### Phase–amplitude coupling maps

Group-level inference on PAC was performed in two stages. First, for each animal, session-level comodulograms were aggregated into a single animal-level observed comodulogram by averaging across sessions. In parallel, an animal-level null distribution was generated by repeatedly resampling one surrogate comodulogram from each session and averaging these across sessions. From this null distribution, the mean and standard deviation at each phase–amplitude frequency pair were estimated, and an animal-level z-scored comodulogram was computed by comparing observed values to the null expectation. Second, group-level inference was performed across animals. For each phase–amplitude frequency pair, the distribution of animal-level z-scores was tested against zero using a one-sided Wilcoxon signed-rank test, assessing whether PAC values were consistently greater than expected under the null. The resulting p-values were corrected for multiple comparisons across the full phase–amplitude frequency grid using the Benjamini–Hochberg false discovery rate (FDR) procedure. The resulting q-value map defines significant regions of PAC at the group level (q < 0.05) and is visualized as −log₁₀(q).

### Histology

After the recording protocol finished, rats were anesthetized with isoflurane (4% induction, 1.5–2% maintenance), and electrolytic lesions (30 µA for 20 s, negative polarity) marked electrode locations. After 48 h of recovery, animals were anesthetized to a surgical plane with ketamine (200 mg/kg) and xylazine (20 mg/kg, intraperitoneally) and euthanized by transcardial perfusion with 0.9% saline followed by 4% paraformaldehyde. Brains were postfixed overnight, transferred to PBS-azide, and sectioned coronally using a vibratome (World Precision Instruments, USA). Sections were Nissl-stained and examined with a Nikon Eclipse CI-L microscope to verify electrode placements.

### Spike Sorting

Extracellular signals were band-pass filtered (600–5,000 Hz), and waveforms with negative or positive peaks crossing a fixed voltage threshold were extracted. Spike sorting was performed with Kilosort2 (https://github.com/MouseLand/Kilosort), and clusters were distinguished using principal-component features.

### Linear mixed-effects models

We used linear and generalized linear mixed-effects models to relate session-level behavioral performance and neural metrics to training and decision strategy. Behavioral performance was quantified as session reward rate, modeled as the number of rewarded trials relative to the total number of valid trials using binomial GLMMs with a logit link. Neural metrics were averaged at the session level and included functional connectivity (weighted phase-lag index, wPLI), directional interactions (spectral Granger causality), and cross-frequency coordination (phase–amplitude coupling, PAC), depending on the specific analysis. Fixed effects included within-animal z-scored training session, behavioral strategy, and, when appropriate, neural predictors or band-limited power covariates. Random effects always included a random intercept for animal, with random slopes for training session added when supported by the data to account for repeated measurements across sessions. Fixed effects were assessed using Type-III tests, and multiple comparisons across neural predictors were controlled using the Benjamini–Hochberg false discovery rate procedure.

### Statistical Analysis

Group differences with a single independent factor were tested with one-way ANOVA when assumptions were met (normality, homogeneity of variance); if normality was violated, we used the Kruskal–Wallis test. For repeated observations, we applied repeated-measures ANOVA when residuals were normally distributed; when only two repeated levels were compared, we used a paired t-test, and if normality was violated we used a Wilcoxon signed-rank test (2 levels) or Friedman test (≥3 levels). Post hoc pairwise comparisons following any omnibus ANOVA were Bonferroni-adjusted. Linear associations between two continuous, normally distributed variables were evaluated with Pearson’s r; otherwise we used Spearman’s ρ. Statistical significance was set at two-tailed α = 0.05. We report effect sizes for all primary tests (Cohen’s d for t-tests, partial η² for ANOVAs, and r/ρ for correlations). All fitting and statistical analyses were performed in MATLAB (MathWorks).

## Data availability

The datasets used and analysed during the current study are available from the corresponding author on reasonable request.

## ACKNOWLEDGMENTS

This manuscript was prepared with the assistance of AI-based tools (ChatGPT and Gemini). AI was used exclusively for improving the clarity, grammar, and overall flow of the text, as well as for providing guidance on statistical analyses, including verification and refinement of the statistical treatment of results, particularly the linear mixed-effects models. In all cases, the authors thoroughly reviewed, edited, and validated any AI-generated content. AI was not used for core research tasks such as experimental design, data acquisition, primary data analysis, interpretation of results, or drawing scientific conclusions. The authors take full responsibility for all content of the manuscript.

## Author contributions

PF designed research and wrote the manuscript; AA performed experiments and data analysis. NE carried out data analysis. GL, MC, GF, and AL-V performed experiments. The authors declare no competing interest.

