## Supplemental information for "Cross-frequency routing in a hippocampo–cortical circuit during probabilistic reversal learning"

**Supplementary Information**

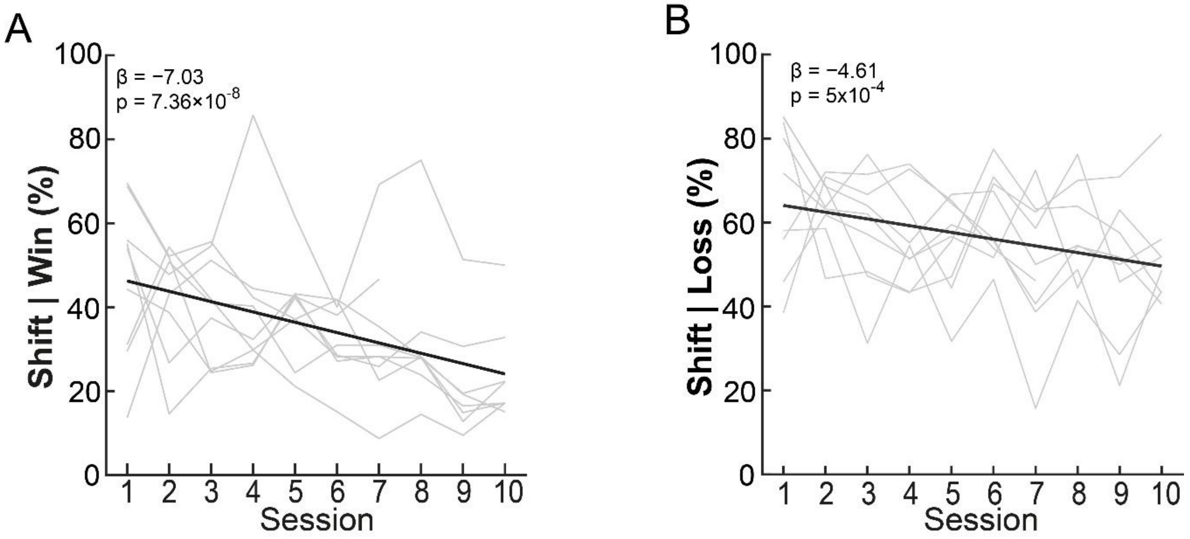


**Figure S1. Outcome-conditioned switching declines across training. A**, Win-shift probability decreased over training ($\beta = -7.03$ , $p = 7.36\times10^{-8}$), indicating reduced exploration after positive outcomes. **B**, The proportion of trials where animals shifted their choice after a single prior loss (computed as 1 – P(stay | 1 prior loss) decreased over training. The mixed-effects model showed a significant negative effect of session ($\beta = -4.61$, $p = 5.26\times10^{-4}$), suggesting a reduced tendency to change choices following isolated negative feedback. Individual trajectories are shown in gray; thick line indicates the fixed-effects prediction.

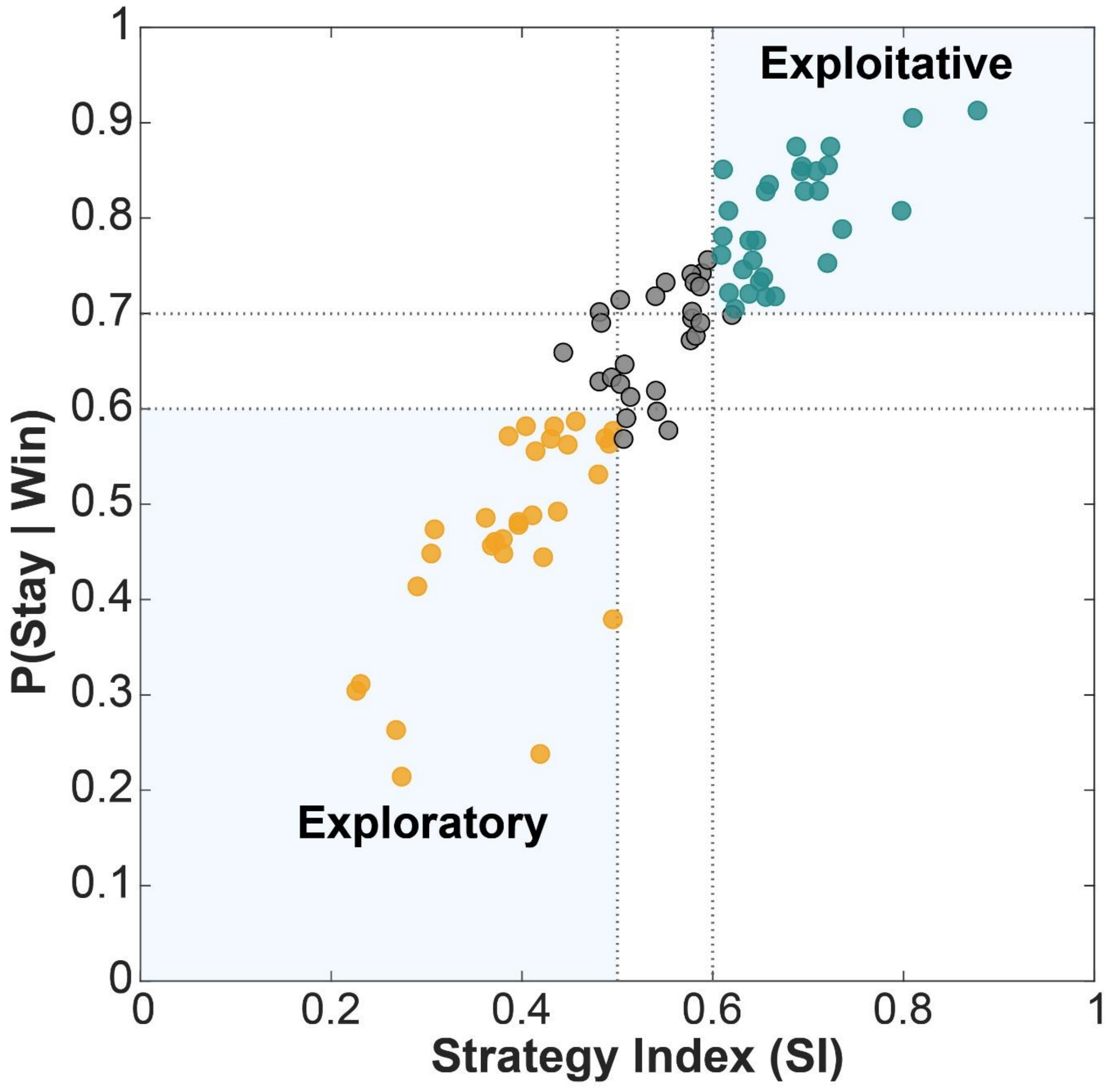


**Figure S2. Strategy classification based on outcome-contingent choice persistence. A**, Strategy plane defined by the probability of staying after a rewarded trial (P (stay|win)) and the Strategy Index (SI = 0.5 x [P(stay|win) + P(stay|loss)]). Each dot represents one session (animal × day). Exploitative (P(stay|win) > 0.70 and SI > 0.60) (n = 30), Exploratory (P(stay|win) < 0.60 and SI< 0.50) (n = 30), and Mixed (intermediate range) (n = 27). Sessions are colored according to their assigned class, and threshold lines denote the decision boundaries.

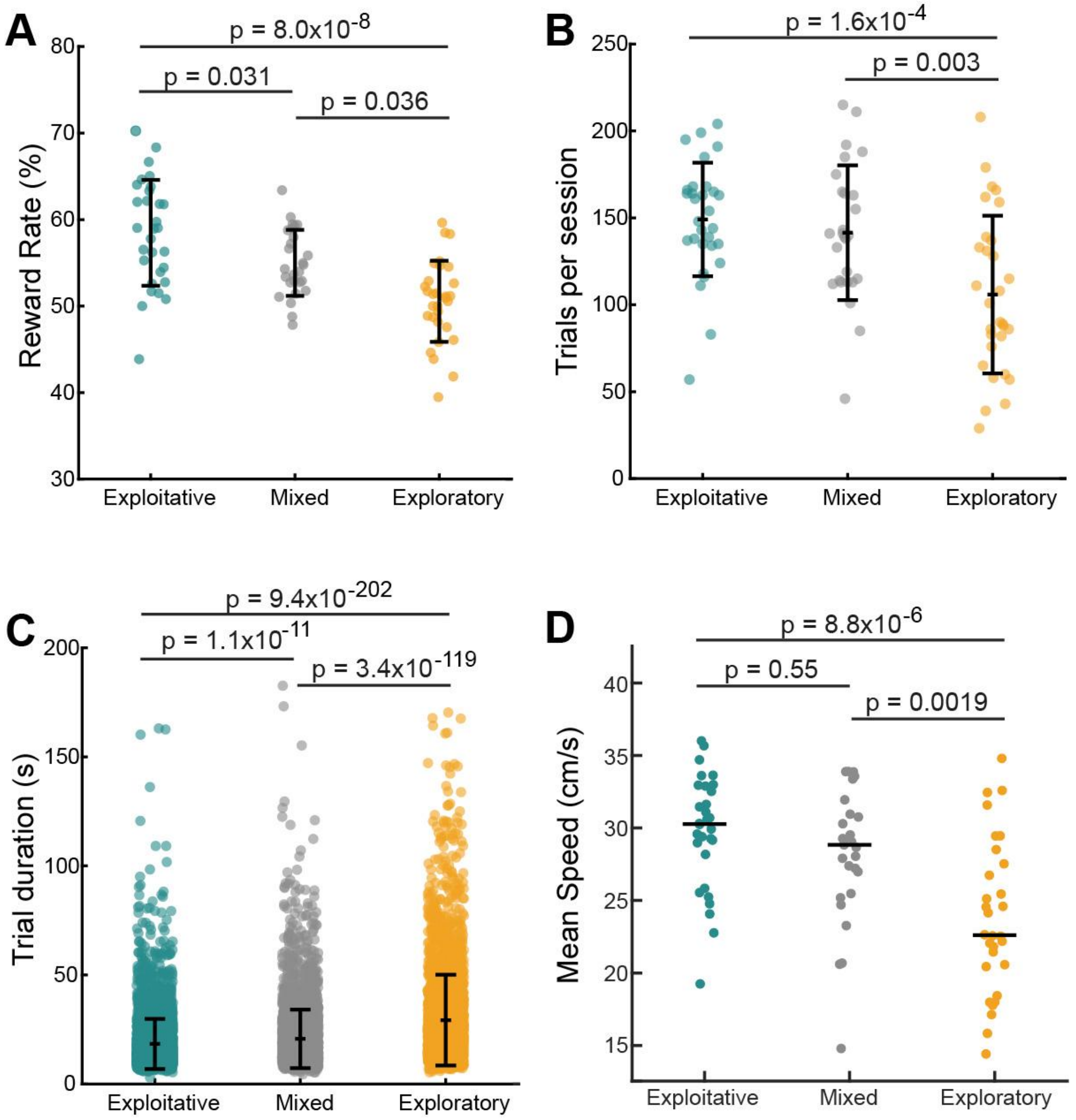


**Figure S3. Behavioral efficiency varies across decision-making strategies. A**, Reward rate plotted for sessions classified as Exploitative, Mixed, or Explorative. Each dot represents one session; error bars indicate group means ± SD. Horizontal significance bars with asterisks denote Bonferroni-corrected post hoc comparisons. A one-way ANOVA revealed a significant main effect of strategy ($F\ (2,84) = 18.92$, $p < 0.0001$). Exploitative sessions showed higher reward rates than Mixed ($p = 0.031$) and Explorative sessions ($p = 8\times10^{-8}$). Mixed sessions also outperformed Explorative sessions ($p = 0.0036$). **B**, A one-way ANOVA revealed significant differences in the number of completed trials across strategies ($p = 10^{-4}$). Exploratory sessions had substantially fewer trials than both Exploitative ($p = 1.6 \times 10^{-4}$) and Mixed sessions ($p = 0.003$). No difference was observed between Exploitative and Mixed sessions ($p = 1$). **C**,

Trial duration differed across strategies ($p = 2.6 \times 10^{-211}$). Exploratory trials were longer than both Mixed ($p = 3.36 \times 10^{-119}$) and Exploitative trials ($p = 9.41 \times 10^{-202}$). Mixed trials were also longer than Exploitative trials ($p = 1.07 \times 10^{-11}$). **D**, Each point is one session; horizontal lines indicate medians. A one-way ANOVA showed a significant effect of strategy ($F(2,84) = 13.36$, $p = 9.2 \times 10^{-6}$). Exploratory sessions were slower than Exploitative ($p = 8.8 \times 10^{-6}$) and Mixed ($p = 0.0019$) sessions, whereas Exploitative and Mixed sessions did not differ ($p = 0.55$).

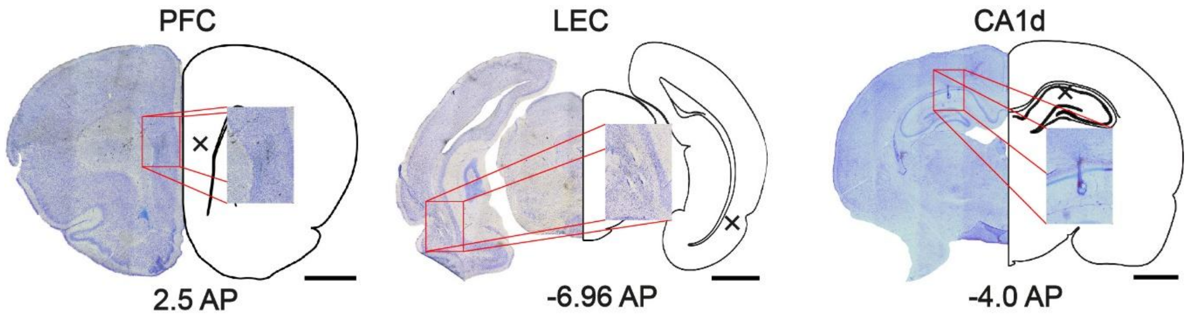


**Figure S4. Histological verification of recording sites.** (PF44; PFC section 56, CA1d section 130, LEC section 167) Example coronal sections showing the location of electrolytic lesions in the PFC, LEC, and CA1d. For each region, the full Nissl-stained section is shown on the left, and a corresponding schematic outline on the right indicates the approximate lesion site (marked with an 'X'). Red boxes denote the area captured in the magnified inset, which highlights the characteristic tissue mark at the lesion core. Scale bar, 2 mm for all panels.

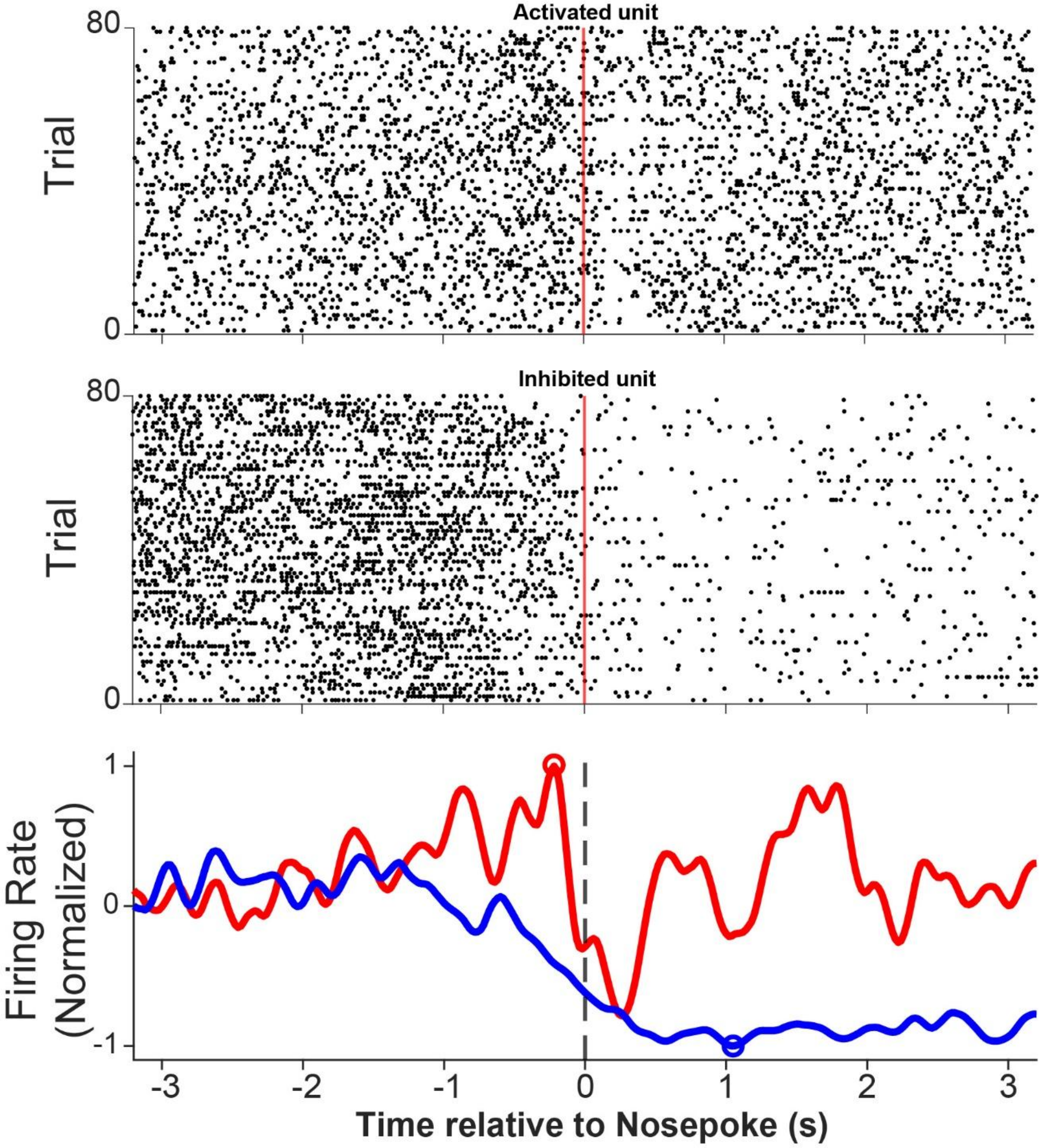


**Figure S5. Example units illustrating the classification of nosepoke-related activation and inhibition.** Top, spike raster of an 'activated' unit aligned to nosepoke (time 0; first 80 valid trials, one row per trial). Middle, spike raster of an 'inhibited' unit recorded in the same session (AA04, Session 1, both PFC units). Middle, same as Top but for an inhibited unit. Bottom, peri-nosepoke normalized firing-rate profiles for the same units shown in Top and Middle (Z-scored to non-nosepoke maze activity and then normalized to each unit's positive or negative peak). Units were classified as activated when $|Z|_{max} > |Z|_{min}$ and as inhibited when $|Z|_{min} > |Z|_{max}$.

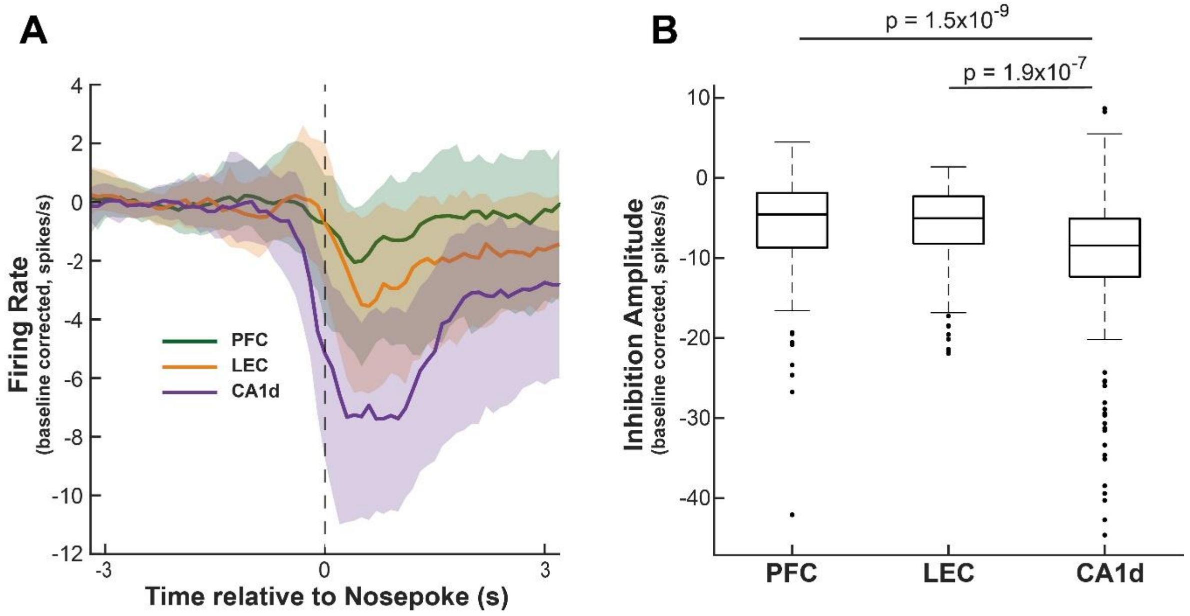


**Figure S6. Peri-goal firing dynamics. A**, Firing rate aligned to the nose-poke (t= 0 s) for PFC, LEC, and CA1d units. Firing rate was computed in real time and corrected by subtracting the trial-by-trial baseline (−3 to −2 s). Curves show the median ± IQR across units. All areas showed a transient suppression, with CA1d exhibiting the largest decrease. **B**, Amplitude of peri-nosepoke inhibition. For each unit, inhibition amplitude was defined as the drop from baseline to the minimum firing rate within 0–2 s after the nosepoke. Distributions differed significantly across areas (Kruskal–Wallis, $p = 5.24 \times 10^{-10}$). Post hoc tests (Mann-Whitney U): PFC vs LEC, $p = 0.398$; PFC vs CA1d, $p = 1.46 \times 10^{-9}$; LEC vs CA1d, $p = 1.94 \times 10^{-7}$.

**Supplementary Tables.**

**Rewards ~ 1 + Session + Trials + Speed + (1 | Animal)**

| Predictor | estimate | SE | t | df | P value | CI lower | CI upper |
|---|---|---|---|---|---|---|---|
| (Intercept) | -0.0409 | 0.0741 | -0.5528 | 83 | 0.5819 | -0.1884 | 0.1064 |
| Trials | 0.002 | 0.0005 | 4.4629 | 83 | 2.5E-05 | 0.0013 | 0.0035 |
| Speed | -1.4403 | 0.4847 | -2.9713 | 83 | 0.0039 | -2.4044 | -0.4762 |
| Session | 0.1470 | 0.0205 | 7.1729 | 83 | 2.8E-10 | 0.1063 | 0.1878 |

**Table S1.** Binomial mixed-effects model of task performance (rewards) including normalized training day (session), the number of valid trials completed in each session (trials), and mean running speed during those trials (speed), with a random intercept for each animal. The training effect (session) remained strongly significant after accounting for these covariates, confirming that improvements in reward rate cannot be attributed to differences in trial count or movement speed across sessions (AIC, 548.93; BIC, 561.26).

**Rewards ~ 1 + Session + Strategy + (1 + Session | Animal)**

| Predictor | estimate | SE | t | df | P value | CI lower | CI upper |
|---|---|---|---|---|---|---|---|
| (Intercept) | 0.0825 | 0.0485 | 1.7008 | 73 | 0.0932 | -0.0142 | 0.1791 |
| Strategy mixed | 0.1047 | 0.0579 | 1.8089 | 73 | 0.0746 | -0.0106 | 0.2201 |
| Strategy exploitative | 0.1839 | 0.0617 | 2.9786 | 73 | 0.0039 | 0.0608 | 0.3069 |
| Session | 0.1261 | 0.0251 | 5.0273 | 73 | 3.4E-06 | 0.0761 | 0.1761 |

| AIC | BIC | Log likelihood | Overdispersion phi |
|---|---|---|---|
| 487.8146 | 504.2212 | -236.9073 | 0.8348 |

| F | df1 | df2 | P value |
|---|---|---|---|
| 4.4415 | 2 | 73 | 0.0151 |

**Table S2**. Binomial GLMM fitted to estimate session-level reward probability and the influence of decision strategy. Model fit statistics (AIC, BIC, log-likelihood) are provided for model comparison. An omnibus Wald test assesses the global effect of decision strategy. The model fit was acceptable (AIC = 487.8) and showed no overdispersion ($\varphi = 0.83$). Adding a training*strategy interaction did not improve model fitness (likelihood-ratio test, $\chi^2 = 4.23$, $P = 0.12$), so we did not include this interaction term in subsequent models.

**A. Rewards ~ 1 + Session + Strategy + Σ [wpli CA1d-LEC+ GC CA1-LEC] + (1 + Session | Animal)**

| Predictor | estimate | SE | t | df | P value | CI lower | CI upper | P FDR |
|---|---|---|---|---|---|---|---|---|
| (Intercept) | 0.2714 | 0.0480 | 5.6484 | 65 | 3.8E-07 | 0.1754 | 0.3674 | - |
| Strategy Exploratory | -0.1586 | 0.0674 | -2.3516 | 65 | 0.0217 | -0.2933 | -0.0239 | - |
| Strategy Mixed | -0.09143 | 0.0558 | -1.6370 | 65 | 0.1064 | -0.2029 | 0.02011 | - |
| Session | 0.1104 | 0.0266 | 4.1556 | 65 | 9.7E-05 | 0.0573 | 0.1635 | - |
| Wpli delta | 0.0173 | 0.0273 | 0.6338 | 65 | 0.5284 | -0.0373 | 0.0719 | 1.000 |
| GC LEC>CA1d delta | -0.0042 | 0.02777 | -0.1502 | 65 | 0.8810 | -0.0596 | 0.0513 | 1.000 |
| GC CA1d>LEC delta | 0.0382 | 0.02606 | 1.4644 | 65 | 0.1479 | -0.0139 | 0.0902 | 1.000 |
| Wpli theta | -0.0015 | 0.0273 | -0.0563 | 65 | 0.9552 | -0.0562 | 0.0531 | 1.000 |
| GC LEC>CA1d theta | 0.0277 | 0.0315 | 0.8783 | 65 | 0.3830 | -0.0353 | 0.0906 | 1.000 |
| GC CA1d>LEC theta | -0.0174 | 0.0327 | -0.5294 | 65 | 0.5984 | -0.0829 | 0.0481 | 1.000 |
| Wpli gamma-slow | -0.0008 | 0.0263 | -0.0287 | 65 | 0.9772 | -0.0533 | 0.0518 | 0.977 |
| GC LEC>CA1d gamma-slow | -0.0023 | 0.0379 | -0.0606 | 65 | 0.9519 | -0.0780 | 0.0734 | 1.000 |
| GC CA1d>LEC gamma-slow | 0.0385 | 0.0280 | 1.3721 | 65 | 0.1748 | -0.0175 | 0.0945 | 1.000 |
| Wpli gamma-fast | -0.0043 | 0.0241 | -0.1793 | 65 | 0.8583 | -0.0526 | 0.0439 | 1.000 |
| GC LEC>CA1d gamma-fast | 0.0089 | 0.0442 | 0.2031 | 65 | 0.8397 | -0.0792 | 0.0972 | 1.000 |
| GC CA1d>LEC gamma-fast | -0.0354 | 0.0364 | -0.9732 | 65 | 0.3340 | -0.1082 | 0.0373 | 1.000 |

| AIC | BIC | Log likelihood | Overdispersion phi | FDR scope | FDR q |
|---|---|---|---|---|---|
| 540.1335 | 585.6280 | -251.0668 | 0.9548 | neural | 0.05 |

**B. Rewards ~ 1 + Session + Strategy + Σ [wpli LEC-PFC+ GC LEC-PFC] + (1 + Session | Animal)**

| Predictor | estimate | SE | t | df | P value | CI lower | CI upper | P FDR |
|---|---|---|---|---|---|---|---|---|
| (Intercept) | 0.2419 | 0.0547 | 4.4195 | 65 | 3.8E-05 | 0.1326 | 0.3512 | - |
| Strategy Exploratory | -0.0918 | 0.0704 | -1.3049 | 65 | 0.1965 | -0.2323 | 0.0487 | - |
| Strategy Mixed | -0.0906 | 0.0555 | -1.6338 | 65 | 0.1071 | -0.2014 | 0.0202 | - |
| Session | 0.1253 | 0.0286 | 4.3840 | 65 | 4.3E-05 | 0.0682 | 0.1824 | - |

| | | | | | | | | |
|---|---|---|---|---|---|---|---|---|
| Wpli delta | 0.0323 | 0.0259 | 1.2491 | 65 | 0.2161 | -0.0194 | 0.0840 | 0.432 |
| GC LEC>PFC delta | 0.01774 | 0.0264 | 0.6714 | 65 | 0.5044 | -0.0350 | 0.0705 | 0.605 |
| GC PFC>LEC delta | -0.0183 | 0.0259 | -0.7034 | 65 | 0.4843 | -0.0702 | 0.0336 | 0.605 |
| Wpli theta | -0.0917 | 0.0286 | -3.2086 | 65 | 0.0021 | -0.1487 | -0.0346 | 0.019 |
| GC LEC>PFC theta | 0.0526 | 0.0278 | 1.8953 | 65 | 0.0625 | -0.0028 | 0.1080 | 0.188 |
| GC PFC>LEC theta | -0.0080 | 0.0228 | -0.3531 | 65 | 0.7252 | -0.0536 | 0.0375 | 0.791 |
| Wpli gamma-slow | 0.01704 | 0.0240 | 0.7092 | 65 | 0.4807 | -0.0309 | 0.0650 | 0.605 |
| GC LEC>PFC gamma-slow | 0.0444 | 0.0354 | 1.2555 | 65 | 0.2138 | -0.0262 | 0.11501 | 0.432 |
| GC PFC>LEC gamma-slow | 0.0054 | 0.0289 | 0.1867 | 65 | 0.8525 | -0.0525 | 0.0633 | 0.852 |
| Wpli gamma-fast | -0.1169 | 0.0381 | -3.0659 | 65 | 0.0032 | -0.1931 | -0.0408 | 0.019 |
| GC LEC>PFC gamma-fast | 0.02818 | 0.0322 | 0.8754 | 65 | 0.3846 | -0.0361 | 0.0925 | 0.605 |
| GC PFC>LEC gamma-fast | 0.0704 | 0.0338 | 2.0806 | 65 | 0.0414 | 0.0028 | 0.1379 | 0.166 |

| **AIC** | **BIC** | **Log likelihood** | **Overdispersion phi** | **FDR scope** | **FDR q** |
|---|---|---|---|---|---|
| 529.3653 | 574.8598 | -245.6826 | 0.6546 | neural | 0.05 |

**C. Rewards ~ 1 + Session + Strategy + Σ [wpli CA1d-PFC+ GC CA1d-PFC] + (1 + Session | Animal)**

| **Predictor** | **estimate** | **SE** | **t** | **df** | **P value** | **CI lower** | **CI upper** | **P FDR** |
|---|---|---|---|---|---|---|---|---|
| (Intercept) | 0.2964 | 0.0445 | 6.6573 | 65 | 6.9E-09 | 0.2075 | 0.3853 | - |
| Strategy exploratory | -0.2131 | 0.0665 | -3.2064 | 65 | 0.0021 | -0.3457 | -0.0804 | - |
| Strategy mixed | -0.1130 | 0.0528 | -2.1415 | 65 | 0.0359 | -0.2184 | -0.0077 | - |
| Session | 0.0932 | 0.0299 | 3.1144 | 65 | 0.0027 | 0.0334 | 0.1529 | - |
| Wpli delta | 0.0576 | 0.0266 | 2.1634 | 65 | 0.0342 | 0.0044 | 0.1108 | 0.410 |
| GC PFC>CA1d delta | -0.0470 | 0.0260 | -1.8070 | 65 | 0.0754 | -0.0989 | 0.0049 | 0.302 |
| GC CA1d>PFC delta | 0.0188 | 0.0258 | 0.7268 | 65 | 0.4699 | -0.0328 | 0.0703 | 0.564 |
| Wpli theta | -0.0108 | 0.0259 | -0.4189 | 65 | 0.6766 | -0.0626 | 0.0408 | 0.738 |
| GC PFC>CA1d theta | 0.0271 | 0.0246 | 1.1009 | 65 | 0.2749 | -0.0220 | 0.0763 | 0.825 |
| GC CA1d>PFC theta | 0.0201 | 0.0233 | 0.8590 | 65 | 0.3934 | -0.0266 | 0.0667 | 0.525 |
| Wpli gamma-slow | 0.0251 | 0.0239 | 1.0474 | 65 | 0.2987 | -0.0227 | 0.0729 | 0.598 |
| GC PFC>CA1d gamma-slow | 0.0572 | 0.0265 | 2.1538 | 65 | 0.0349 | 0.0041 | 0.1102 | 0.210 |

| | | | | | | | | |
|---|---|---|---|---|---|---|---|---|
| GC CA1d>PFC gamma-slow | 0.0265 | 0.0270 | 0.9827 | 65 | 0.3293 | -0.0273 | 0.0804 | 0.494 |
| Wpli gamma-fast | -0.02495 | 0.0243 | -1.0257 | 65 | 0.3088 | -0.0735 | 0.0236 | 0.529 |
| GC PFC>CA1d gamma-fast | -0.0279 | 0.0255 | -1.0925 | 65 | 0.2786 | -0.0790 | 0.0231 | 0.669 |
| GC CA1d>PFC gamma-fast | -0.0095 | 0.0261 | -0.3658 | 65 | 0.7156 | -0.0616 | 0.0425 | 0.716 |

| **AIC** | **BIC** | **Log likelihood** | **Overdispersion phi** | **FDR scope** | **FDR q** |
|---|---|---|---|---|---|
| 534.0533 | 579.5478 | -248.0266 | 0.8708 | neural | 0.05 |

**Table S3.** Mixed-effects model linking reward rate to behavioral strategy and neural connectivity for the recorded pairs of regions: **A**, CA1d-LEC; **B**, LEC-PFC; **C**, CA1d-PFC. Generalized linear mixed-effects model (GLMM) predicting session-level reward rate from behavioral strategy, within-rat normalized training day (session), and residualized neural connectivity metrics (WPLI and bidirectional Granger Causality) computed across delta, theta, slow-gamma, and fast-gamma bands. Connectivity terms were residualized against band-limited power in both areas to isolate effects beyond spectral amplitude. The model includes random intercepts and session slopes for each animal. Fixed-effect estimates, standard errors, raw P-values, and FDR-adjusted P-values (Benjamini–Hochberg, q = 0.05) are reported. Model fit statistics (AIC, BIC, and log-likelihood) are provided to assess overall model fit and enable comparison across model specifications. An omnibus Wald F test evaluates the global contribution of multi-level predictors (e.g., behavioral strategy).

**A. Log(PAC CA1d-PFC) ~ 1 + Session + Strategy + [z-logP theta-phase + z-logP gamma-fast amplitude] + (1 + Session | Animal)**

| Predictor | estimate | SE | t | df | P value | CI lower | CI upper |
|---|---|---|---|---|---|---|---|
| (Intercept) | -16.7898 | 3.6601 | -4.5872 | 75 | 1.7E-05 | -24.0812 | -9.4985 |
| Strategy Exploratory | 0.0750 | 0.1464 | 0.5125 | 75 | 0.6097 | -0.2167 | 0.3669 |
| Strategy Mixed | 0.0716 | 0.1159 | 0.6176 | 75 | 0.5386 | -0.1594 | 0.3027 |
| Session | -0.0281 | 0.0933 | -0.3007 | 75 | 0.7644 | -0.2139 | 0.1578 |
| z-logP CA1d theta PAC | 0.0057 | 0.0571 | 0.1008 | 75 | 0.9199 | -0.1080 | 0.1195 |
| z-logP PFC gamma-fast PAC | -0.0102 | 0.0696 | -0.1468 | 75 | 0.8836 | -0.1489 | 0.1284 |

| Observations | AIC | BIC | Log likelihood | Residual Std |
|---|---|---|---|---|
| 81 | 172.3844 | 195.5593 | -76.1922 | 0.3452 |

**B. Log(PAC CA1d-LEC) ~ 1 + Session + Strategy + [z-logP theta-phase + z-logP gamma-fast amplitude] + (1 + Session | Animal)**

| Predictor | estimate | SE | t | df | P value | CI lower | CI upper |
|---|---|---|---|---|---|---|---|
| (Intercept) | -16.2872 | 3.7331 | -4.3628 | 75 | 4.0E-05 | -23.7239 | -8.8504 |
| Strategy Exploratory | -0.0492 | 0.0789 | -0.6237 | 75 | 0.5347 | -0.2065 | 0.1080 |
| Strategy Mixed | 0.0794 | 0.0593 | 1.3386 | 75 | 0.1847 | -0.0387 | 0.1977 |
| Session | -0.0448 | 0.0320 | -1.3973 | 75 | 0.1664 | -0.1087 | 0.0190 |
| z-logP CA1d theta PAC | 0.0384 | 0.0285 | 1.3480 | 75 | 0.1816 | -0.0183 | 0.0952 |
| z-logP LEC gamma-fast PAC | 0.0455 | 0.0239 | 1.900 | 75 | 0.0612 | -0.0022 | 0.0932 |

| Observations | AIC | BIC | Log likelihood | Residual Std |
|---|---|---|---|---|
| 81 | 95.2196 | 118.3944 | -37.6098 | 0.2007 |

**Table S4.** Linear mixed-effects models relating hippocampal theta–phase to cortical gamma–amplitude coupling to training session and behavioral strategy. **A**, log(PAC) CA1d-PFC and **B**, log(PAC) CA1d-LEC. Each row represents a fixed-effect term from a model fitted on Animal × Session. The dependent variable is the log-transformed PAC modulation index within a CA1d θ (5–10 Hz) × cortical fast-γ (60–110 Hz). Fixed effects include z-scored training session (Session), strategy (Exploitative, Mixed, Exploratory), and within-rat z-scored θ power in CA1d and fast-γ power in the corresponding cortical area. Both models include random intercepts and random slopes for Session by animal (1 + Session|Animal). Model-specific fit statistics (number of observations, AIC, BIC, log-likelihood, and residual standard deviation) are reported separately for panels A and B to characterize overall model fit.

**Rewards ~ 1 + Session + WPLI + PAC CA1d-PFC + PAC CA1d-LEC + PAC LEC-PFC + PAC PFC-LEC + (1 + Session | Animal)**

| Predictor | estimate | SE | t | df | P value | CI lower | CI upper |
|---|---|---|---|---|---|---|---|
| (Intercept) | 0.2952 | 0.0421 | 7.0024 | 67 | 1.4E-09 | 0.2110 | 0.3793 |
| Strategy exploratory | -0.2184 | 0.0634 | -3.4444 | 67 | 0.0009 | -0.3450 | -0.0918 |
| Strategy mixed | -0.0977 | 0.0504 | -1.9387 | 67 | 0.0567 | -0.1984 | 0.0028 |
| Session | 0.1148 | 0.0267 | 4.2873 | 67 | 5.9E-05 | 0.0613 | 0.1682 |
| Wpli LEC-PFC theta | -0.0430 | 0.0243 | -1.7703 | 67 | 0.0812 | -0.0916 | 0.0054 |
| Wpli LEC-PFC gamma-fast | -0.0328 | 0.0249 | -1.3166 | 67 | 0.1924 | -0.0825 | 0.0169 |
| Wpli PFC-CA1d theta | -0.0092 | 0.0255 | -0.3617 | 67 | 0.7186 | -0.0603 | 0.0418 |
| Wpli PFC-CA1d gamma-fast | -0.0097 | 0.0236 | -0.4129 | 67 | 0.6809 | -0.0570 | 0.0374 |
| Wpli LEC-CA1d theta | -0.0033 | 0.0222 | -0.1494 | 67 | 0.8816 | -0.0477 | 0.0411 |
| Wpli LEC-CA1d gamma-fast | -0.0023 | 0.0242 | -0.0956 | 67 | 0.9240 | -0.0508 | 0.0461 |
| PAC CA1d-PFC | -0.0334 | 0.0260 | -1.2848 | 67 | 0.2032 | -0.0853 | 0.0185 |
| PAC CA1d-LEC | -0.0115 | 0.0246 | -0.4685 | 67 | 0.6409 | -0.0607 | 0.0376 |
| PAC LEC-PFC | 0.0043 | 0.0224 | 0.1952 | 67 | 0.8457 | -0.0404 | 0.0492 |
| PAC PFC-LEC | 0.0221 | 0.0231 | 0.9602 | 67 | 0.3403 | -0.0239 | 0.0683 |

**Table S5.** Session-level generalized linear mixed-effects model (binomial, logit link) relating reward rate to training day (session), behavioral strategy, and residualized coupling metrics. The dependent variable was the number of rewarded trials per session (Rewards) with the total number of valid trials (Trials) as binomial denominator. Fixed effects included within-rat z-scored training session (Session). Strategy (Exploitative, Mixed, Exploratory; reference category as in the table), residualized WPLI for three area pairs (LEC–PFC, PFC–CA1d, LEC–CA1d) in theta (5–11 Hz) and fast-gamma (60–110 Hz), and residualized PAC in four directions (CA1d→PFC, CA1d→LEC, LEC→PFC, PFC→LEC). WPLI and PAC predictors were z-scored residuals obtained after regressing each measure, within each animal, on the corresponding band-limited power in the contributing areas.

**log(GC LEC>CA1d gamma-slow) ~ 1 + Session + Strategy + z-logP LEC gamma-slow + z-logP CA1d gamma-slow + (1 + Session | Animal)**

| Predictor | estimate | SE | t | df | P value | CI lower | CI upper |
|---|---|---|---|---|---|---|---|
| (Intercept) | -6.9920 | 3.6250 | -1.9288 | 75 | 0.0575 | -14.2136 | 0.2294 |
| Strategy Exploratory | -0.0685 | 0.0577 | -1.1871 | 75 | 0.2389 | -0.1835 | 0.0464 |
| Strategy Mixed | -0.0456 | 0.0474 | -0.9623 | 75 | 0.3389 | -0.1401 | 0.0488 |
| Session | 0.1071 | 0.0341 | 3.1347 | 75 | 0.0024 | 0.0391 | 0.1752 |
| z-logP LEC gamma-slow | -0.0227 | 0.0186 | -1.2183 | 75 | 0.2268 | -0.0597 | 0.0143 |
| z-logP CA1d gamma-slow | 0.0112 | 0.0263 | 0.4207 | 75 | 0.6751 | -0.0414 | 0.0635 |

| Predictor | F | df1 | df2 | P value |
|---|---|---|---|---|
| (Intercept) | 3.7202 | 1 | 75 | 0.0575 |
| Strategy | 0.7811 | 2 | 75 | 0.4615 |
| Session | 9.8268 | 1 | 75 | 0.0024 |
| z logP LEC gamma-slow | 1.4844 | 1 | 75 | 0.2268 |
| z logP CA1d gamma-slow | 0.1770 | 1 | 75 | 0.6751 |

**log(GC CA1d>LEC gamma-slow) ~ 1 + Session + Strategy + z-logP LEC gamma-slow + z-logP CA1d gamma-slow + (1 + Session | Animal)**

| Predictor | estimate | SE | t | df | P value | CI lower | CI upper |
|---|---|---|---|---|---|---|---|
| (Intercept) | -7.8858 | 3.5418 | -2.2264 | 75 | 0.0289 | -14.9416 | -0.8300 |
| Strategy Exploratory | 0.0425 | 0.0606 | 0.7019 | 75 | 0.4849 | -0.0782 | 0.1633 |
| Strategy Mixed | -0.0005 | 0.0507 | -0.0098 | 75 | 0.9921 | -0.1016 | 0.1006 |
| Session | 0.1190 | 0.0470 | 2.5270 | 75 | 0.0136 | 0.0251 | 0.2128 |
| z logP LEC gamma-slow | -0.0234 | 0.0198 | -1.1823 | 75 | 0.2407 | -0.0629 | 0.0160 |
| z logP CA1d gamma-slow | 0.0418 | 0.0309 | 1.3491 | 75 | 0.1813 | -0.0199 | 0.1035 |

| Predictor | F | df1 | df2 | P value |
|---|---|---|---|---|
| (Intercept) | 4.957073491 | 1 | 75 | 0.028986554 |
| Strategy | 0.364568215 | 2 | 75 | 0.695720358 |
| Session | 6.386201376 | 1 | 75 | 0.01360739 |
| z logP LEC gamma-slow | 1.398041054 | 1 | 75 | 0.240786858 |
| z logP CA1d gamma-slow | 1.820261201 | 1 | 75 | 0.181341304 |

**Table S6.** Mixed-effects models of gamma-slow Granger causality between LEC and CA1d. Fixed-effects results of linear mixed-effects models predicting gamma-slow (30–60 Hz) Granger causality for CA1d→LEC **(A)** and LEC→CA1d **(B)**. Each model included session (z-scored within rat), Strategy, and band-limited power in LEC and CA1d as fixed effects, with random intercepts and Session slopes per animal. Significance of fixed effects was assessed using marginal Wald ANOVA tests with residual degrees of freedom (DFMethod = 'Residual'), reporting F statistics, numerator/denominator degrees of freedom, and p-values.
